# Uncovering cancer dependencies in peptide-interacting protein pockets

**DOI:** 10.64898/2026.03.13.711608

**Authors:** Mihkel Örd, Liz Kogan, Devin Bradburn, Michelle Leiser, Mardo Kõivomägi, Norman E. Davey, Pau Creixell

## Abstract

Cancer cells often become dependent on specific molecular functions. As many proteins perform multiple functions mediated by different pockets and interfaces, we hypothesized that we could identify distinct cancer dependencies and therapeutic vulnerabilities by disrupting peptide-binding pockets. To test this hypothesis, we screened a proteome-wide library of 7152 genetically encoded peptides across nine cancer cell lines. We identify common and selective dependencies on peptide-binding pockets and find that gene knockout and peptide-mediated inhibition of pockets often drive divergent phenotypes. For the common-essential gene HCF1, we identify a therapeutic window by using inhibitory peptides with varying affinity. Moreover, peptides targeting TLE1-4 reveal a dependency hidden in genetic screens by homolog redundancy. We also uncover that peptides inhibiting cyclin D drive specific suppression of leukemia cell proliferation and demonstrate that these peptides improve the potency of CDK4/6 inhibitors. Overall, our screening platform facilitates data-driven prioritization of molecular pockets for subsequent therapeutic translation.

## Introduction

As they evolve and acquire malignant phenotypes, cancer cells often become dependent on specific genes for survival and proliferation^1,2^. These differential dependencies lay the basis for targeted cancer therapies that aim to specifically kill cancer cells while minimizing toxicity to other cells. For example, the development of tyrosine kinase inhibitors targeting the oncogenic fusion BCR-ABL has revolutionized the treatment of chronic myeloid leukemia (reviewed in^3^), and CDK4/6 inhibitors are used in advanced breast cancer based on the principle that these cancer cells become addicted to increased kinase activity^4^. These successes have motivated genome-wide genetic screens to identify specific dependencies of distinct cancers and to inform on target prioritization for drug design^5,6^.

While the development of CRISPR-based technologies has accelerated the identification of cancer gene dependencies even further, with “dependency maps” already available for over a thousand cell lines^5^, many CRISPR dependencies remain undruggable and additional dependencies may have been missed due to genetic redundancies. Moreover, the effect of inhibiting a pocket may differ from the effect of knocking the gene out or down. For example, inhibiting cyclin B protein interactions leads to a cancer-specific anti-proliferative effect driven by specific de-repression of a negative regulator whose binding is outcompeted by the drug^7^.

In parallel, screens characterizing dependencies on specific protein-protein interactions^8^ and identification of anti-proliferative peptides and proteins have more recently emerged to provide information on the phenotype of inhibiting specific interactions or pockets, thereby providing a direct molecular interface to target or even a potential peptide/protein-based therapeutic^9,10^. Supporting the therapeutic relevance of these approaches, recent years have brought an increasing number of small molecule compounds that effectively disrupt protein-protein interactions^11,12^, increasing our existing cancer therapy arsenal beyond inhibitors of catalytic sites and extracellular receptors. A classic example for the therapeutic value of targeting peptide-binding pockets is the inhibition of Bcl-2, a pro-survival protein that when binding to pro-apoptotic proteins via BH3 motifs inhibits apoptosis. This interaction is outcompeted by venetoclax, a drug that is used in the treatment of leukemia^13^, and that elegantly showcases the potential of targeting peptide-mediated interactions.

Human cells are estimated to rely on over 100000 protein-protein interactions mediated by short linear motifs^14^. Unlike domain-domain protein-protein interactions, which rely on large surfaces, peptide-mediated interactions, where a 3-12 amino acid long peptide motif binds to a pocket on a folded domain^14^, typically offer more accessible and druggable pockets that can be exploited for therapeutic benefit. While peptide-mediated interactions are often dynamic and low affinity, they control key aspects of protein function including localization, stability, modification state and binding partners. They are frequently present in signaling hub proteins that interact with a large number of diverse proteins via convergently evolved motifs. Yet, despite the significant potential of targeting them, less than 10% of the expected 100000 peptides that mediate protein-protein interactions within the human proteome have been experimentally characterized so far, and as a result the phenotype and therapeutic potential of inhibiting the vast majority of peptide-binding pockets is currently unknown^14,15^. The findings that simultaneous targeting of multiple pockets in different or even the same target proteins enable new and attractive therapeutic opportunities further motivates our systematic study of peptide-binding pockets.

Thus, we here investigate the phenotype of inhibiting specific peptide-protein interactions at a proteome-wide scale in nine cancer cell lines by performing pooled genetically expressed competitor peptide dropout screens (**Fig. 1A**). We find that while some peptide binding pockets are common essential, the phenotype of inhibiting some others is cell line or cancer-type specific. We explore the mechanisms driving the peptide phenotype focusing on three peptide classes, namely HCF1, TLE1-4, and cyclin D1-3 targeting peptides. By demonstrating that the sensitivity to competitor peptides does not correlate with target genetic dependency, we characterize the added value that screening pocket inhibitors can provide for target prioritization.

**Figure 1.**
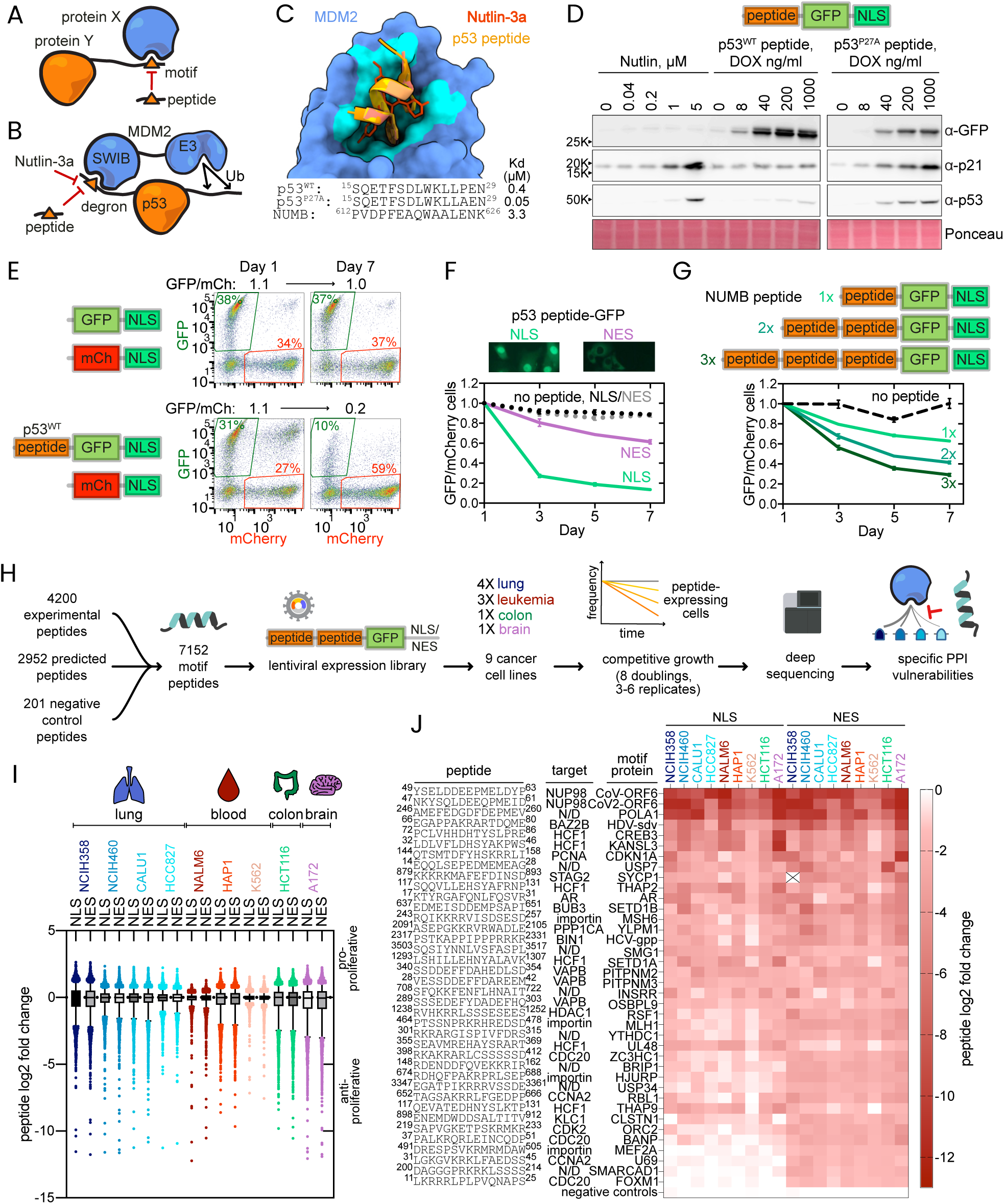
Pooled competitor peptide screening identifies dependencies in peptide-binding pockets. (**A**) Scheme showing how an overexpressed motif peptide from protein Y could outcompete the endogenous interaction between proteins X and Y and other interactions involving that pocket on protein X. (**B**) MDM2 SWIB domain binds to a degron motif in p53, leading to p53 degradation and maintenance of low p53 expression. (**C**) Superimposed structures of MDM2 SWIB domain in complex with the p53 degron peptide and Nutlin-3a. (**D**) Western blot images showing the dose-response of HCT116 cells to Nutlin-3a and p53 degron competitor peptides. (**E**) Flow cytometry plots showing the change in GFP-positive and mCherry-positive cell fractions in a pairwise competitive growth experiment in HCT116 cells. (**F**) Fusion with NLS and NES motifs localises the competitor peptides to different subcellular compartments, which impacts the anti-proliferative effect of the p53 peptide, as observed in a pairwise GFP/mCherry cell competitive growth experiment in HCT116. (**G**) Pairwise competitive growth experiments showing the impact of motif repeats on the potency of competitors based on NUMB MDM2 degron in HCT116. (**H**) Experimental scheme of proteome-wide competitor peptide screening to identify peptide-binding pocket inhibition vulnerabilities. (**I**) The distribution of peptide abundance changes in 9 cancer cell lines. The error bars show 5-95% percentile. (**J**) Heatmap showing dropout scores (log2 fold change) of the peptides identified as anti-proliferative in at least 7 cell lines.

## Results

### Expression of MDM2 competitor peptide can mirror Nutlin-3a effect

To optimize and test the potential of screening genetically encoded competitor peptide libraries, we chose the SWIB domain of MDM2 as our target model system for initial testing. MDM2 is an E3 ubiquitin ligase that suppresses p53 activity by mediating its proteasomal degradation through an interaction that can be inhibited by the small molecule drug Nutlin-3a **(Fig. 1B,C)**. To compare the efficacy of inhibiting the MDM2-p53 interaction with genetically encoded peptides and Nutlin-3a, we expressed a doxycycline-inducible p53 degron peptide fused to EGFP in HCT116, a colorectal adenocarcinoma cell line harboring a functional p53 signaling pathway **(Fig. 1D)**. We observed that the high-affinity competitor p53^P27A^ peptide (a 15-amino-acid long peptide derived from p53 with a P27A mutation that increases its MDM2 affinity)^16^ and Nutlin-3a caused a similar dose-dependent stabilization of p53 and increased expression of its downstream target p21 (**Fig. 1D**). The finding that the wild-type p53 peptide triggered a reduced p21 and p53 upregulation **(Fig. 1D)** illustrates how peptides with varying affinities or specificities may elicit different molecular and phenotypic outcomes.

Next, we studied the effect of the MDM2-targeting peptide on proliferation. Using a pairwise competitive growth experiment coupled with flow cytometry we found that, unlike no peptide controls, the p53-MDM2 competitor peptide inhibited proliferation of p53-wild-type HCT116 cells, but not of p53 mutant HCC827 cells **(Fig. 1E, S1A)**. These experiments suggest that genetically expressed motif competitor peptides can recapitulate small molecule drug effects, which led us to hypothesize that they could be used in pooled competitive growth dropout screens to investigate the phenotype of inhibiting peptide-binding pockets proteome-wide.

### Localization signals and motif repeats can improve the potency of competitor peptides

Given that the majority of known and estimated binding affinities of the peptide-mediated protein-protein interactions are in the low micromolar range **(Fig. S1B)**^17^, we wondered whether we could increase the sensitivity of our screens by optimizing where and how our competitor peptides are expressed. First, we tested the impact of compartmentalizing the peptide to either the nucleus or the cytoplasm, by adding nuclear localization (NLS) and nuclear export signals (NES), respectively **(Fig. 1F).** We observed that our NLS construct led to a stronger anti-proliferative effect of the p53 MDM2-targeting peptide than its NES counterpart **(Fig. 1F)**, most likely by increasing the competitor peptide concentration in the nucleus where MDM2 is localized^18^. In addition to increasing the sensitivity of our screens, we reasoned that the ability to screen and score peptide pools in both NLS and NES configurations may provide added value by narrowing down the mechanism of action and localization of the major target of our competitor peptides.

As an orthogonal strategy to increase the sensitivity of our screens we investigated the impact of increasing the number of repeats of each peptide in our competitor constructs, which had been previously found to improve the potency of a PP2A inhibitor and the ability of peptides pulling down their targets^19,20^. While increasing the number of peptide repeats did not affect the anti-proliferative effect of the high-affinity p53 peptide (**Fig. S1C**), we found that the inhibitory effect of a ten-times lower affinity NUMB peptide was improved by increasing the number of peptide repeats **(Fig. 1G)**. These results suggest that introducing multiple repeats of the motif into the competitor scaffold can improve the effect of weaker-affinity peptides. Thus, by incorporating localization signals and multiple peptide repeats, we improved the sensitivity of our screens to identify molecular pocket dependencies.

### Pooled proteome-wide dropout screening identifies anti-proliferative competitor peptides

We next performed a pooled peptide dropout screen in cancer cell lines to identify common, cancer-type- and cell-line-specific pocket dependencies covering peptide-domain interactions proteome-wide. To this end, we assembled a library of 7152 fifteen-residue long peptide sequences including 4200 experimentally validated peptides, 2952 predicted motif-containing peptides, and 201 negative control peptides **(Fig. 1H, S1D-E)**. As negative controls, we included 100 scrambled peptides and 101 mutant peptides, where key residues mediating major peptide-domain interactions were mutated to alanine. We investigated the effect of our peptide pool on the proliferation of 4 lung cancer cell lines (NCIH358, NCIH460, CALU1, HCC827), three leukemia cell lines (HAP1, NALM6, K562), one colon cancer cell line (HCT116), and one brain cancer cell line (A172) by performing lentiviral transductions followed by competitive growth experiments **(Fig. S1F).** We found that up to 9.6% of the peptides in our library were anti-proliferative in each cell line (**Fig. 1I, Fig. S1G**). Most anti-proliferative peptides inhibited either a single or a small number of cell lines, while 92 peptides (7.7% of all hits) were anti-proliferative in at least six out of the nine cancer cell lines (**Fig. S1H**). The anti-proliferative peptides were of diverse origin (including docking interaction and substrate peptides) and were known or predicted to target a variety of protein domains (**Fig. S1I-J**). There was no difference in the fraction of anti-proliferative peptides between experimentally characterized and predicted binding peptides (**Fig. S1K**), suggesting that a significant fraction of the predicted motif peptides indeed bind and inhibit protein domains. We found no considerable correlation between general biophysical characteristics (charge and hydrophobicity) of our peptides and their anti-proliferative activity, further supporting that they likely operate through specific targeting rather than general cytotoxicity (**Fig. S1L-M**).

The most anti-proliferative peptides included NUP98/RAE1-targeting peptides from SARS-CoV1/2 ORF6, a protein that inhibits nuclear export of mRNAs and shuttling of proteins^21^, a PCNA-targeting peptide from CDKN1A, and multiple Host Cell Factor 1 (HCF1) binding peptides among others (**Fig. 1J**). CDKN1A (p21) is known to inhibit DNA replication by sequestering PCNA^22^, and, alanine mutations that disrupt the PCNA binding motif in the CDKN1A and ScUNG1 peptides also decrease the toxicity of these peptides (**Fig. S1N**). The phenotype of inhibiting HCF1 interactions, however, had not been characterized, motivating HCF1 to become our first interaction of interest as we further discuss in the next sections. Altogether, our proteome-wide competitor peptide screen identified anti-proliferative peptides as well as phenotypic readouts of inhibiting different interaction pockets in a range of cancer cell lines.

### Identifying common essential and selectively essential peptide-binding pocket dependencies

To facilitate analysis, we next grouped all peptides that contain the same motif and thus are expected to target the same pocket and investigated the behavior of these peptide sets. Given that there was minimal difference between the anti-proliferative effect of all targeting peptides taken together and our scrambled and non-targeting negative controls (**Fig. 2A**), we concluded that the majority of competitor peptides do not significantly impact cell proliferation. However, specific groups of peptides showed significant anti-proliferative effects across all or a subset of the nine cell lines tested. For example, 8 peptides targeting the BET domain of bromodomain proteins (BET-BRD) were among the anti-proliferative peptides in NALM6 (p-value 3.03*10^-6^), as were the majority of the 25 HCF1-targeting peptides in the library (p-value 4.15*10^-10^) (**Fig. 2A**), suggesting that this cell line is dependent on these peptide-binding pockets. Conversely, a small number of peptide sets increased proliferation, including peptides targeting the ankyrin domain of ANKRA2 and RFXANK (**Fig. 2A**), two tumor suppressors that are activated by p53^23^.

**Figure 2.**
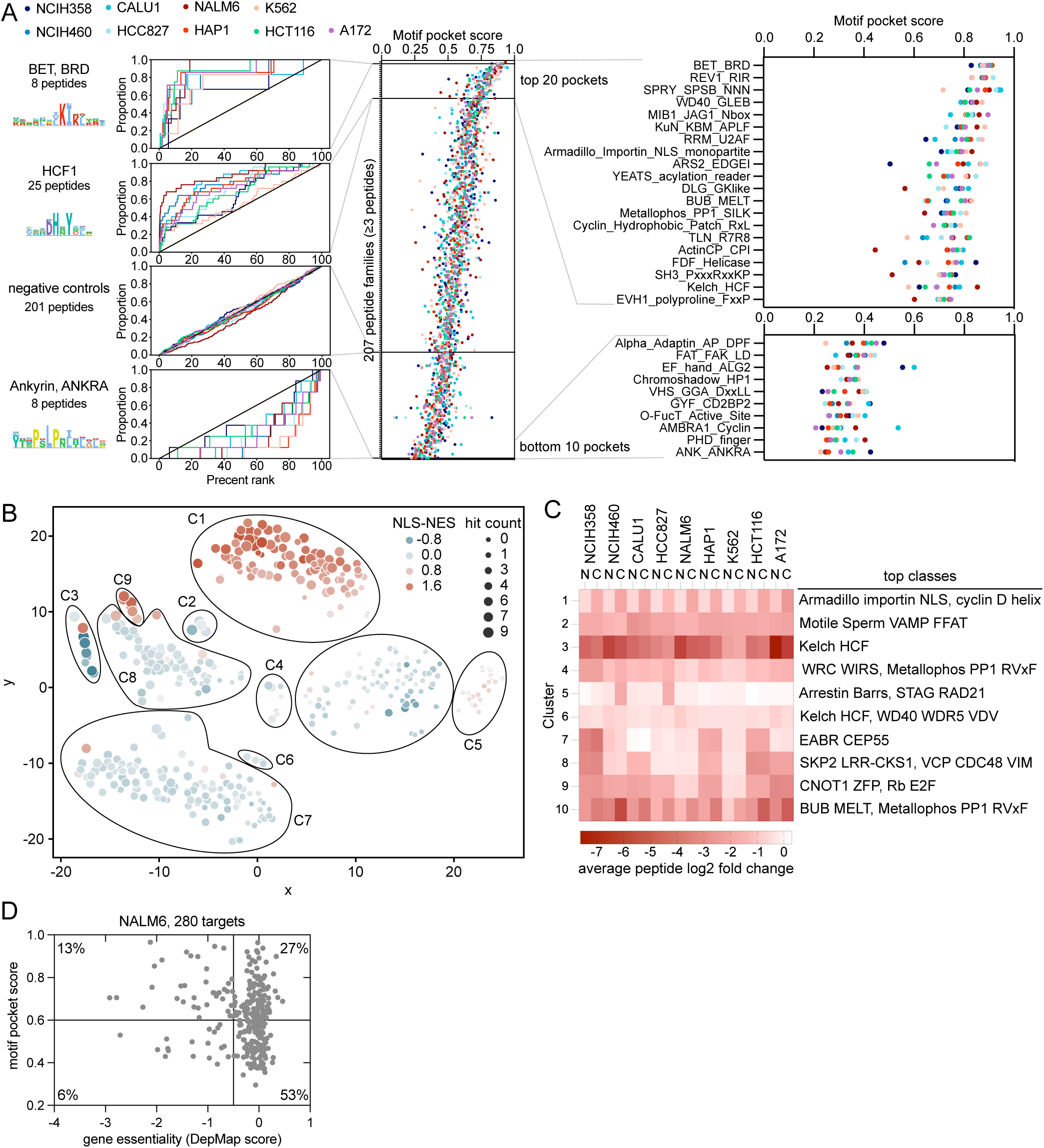
Cancer cell lines exhibit varying sensitivities to inhibition of peptide-binding pockets. (**A**) Left: cumulative rank plots of peptides from the indicated peptide groups in 9 cell lines in the proteome-wide dropout screen. Middle and right: plots showing the motif pocket scores, calculated as the area under the cumulative rank plots, of all motif families with at least 3 peptides. (**B**) tSNE dimensionality reduction shows the distribution and HDBSCAN clustering of the top 2% anti-proliferative peptides detected in the screen from all cell lines. (**C**) Heatmap showing the average dropout score of peptides from the different clusters and the top peptide classes based on the enrichment in each cluster. (**D**) Scatterplot showing the motif pocket scores calculated as in panel ‘A’ and the gene essentiality scores of the target proteins. The gene essentiality data was obtained from DepMap CRISPR knockout screens.

Among the families of anti-proliferative peptides, some are similarly anti-proliferative in all tested cell lines, suggesting that they inhibit common essential interactions. These include peptides targeting BET domains and the REV1 C-terminal domain (**Fig. 2A, S2A**), the latter also being the target of small molecule REV1 inhibitors that sensitize cancer cells to DNA damage therapies^24^. In contrast, other peptide sets show significant variation among cell lines, including HCF1 binding peptides, which rank among the most anti-proliferative in NALM6, while their effect in HCT116 and NCIH358 is considerably reduced (**Fig. 2A**). From the 7152 peptides in our library, we were able to collapse 3810 of them into 206 motif classes containing at least 3 peptides (**Fig. S2B**). This proteome-wide analysis showed us that a small number of peptide sets inhibit common essential interaction pockets, while a larger number of them uncover more specific vulnerabilities. Through this analysis, we identified candidate cancer-type-specific vulnerabilities, including cyclin D substrate docking peptides, which significantly inhibited all three leukemia cell lines in our panel while showing little effect in solid cancer cell lines (**Fig. S2C**). For this reason, we selected cyclin D peptides as our second interaction of interest as we show in the next sections. Thus, altogether, our competitor peptide screening technology may facilitate the identification of both common essential and more selective interaction pocket vulnerabilities.

Investigating the top 2% most anti-proliferative peptides from each cell line, we found that the peptides clustered largely by the cell lines they inhibited and the effect of nuclear or cytoplasmic compartmentalization (NLS- or NES-fused scores) (**Fig. 2B**). For example, cluster 1 consists of peptides that were strongly anti-proliferative in most of the tested cell lines and shared a stronger effect when fused to a NES, whereas cluster 3 contained similarly anti-proliferative peptides that showed a stronger effect when fused to a NLS or that showed little differences between localization signals (**Fig. 2B-C**). Some clusters were significantly enriched in specific peptide classes, such as cluster 3, where 8 out of the 13 peptides were HCF1-targeting peptides (p-value 1.35*10^-14^), and cluster 1, which was dominated by “Armadillo Importin NLS” motifs (p-value 1.15*10^-24^) (**Fig. 2C**). We also identified clusters of peptides affecting different subsets of cell lines, such as cluster 4 that has the strongest effect in NCIH358, and cluster 8 with peptides most anti-proliferative in five of the tested lines (**Fig. 2C**). Thus, our proteome-wide peptide screen allows us to identify diverse patterns of cell line vulnerabilities upon inhibition of specific protein-protein interactions.

While essential proteins are more likely to contain an essential pocket (p-value < 0.00001, chi-squared test), we did not observe a correlation between CRISPR-based gene essentiality and peptide-based pocket essentiality obtained from the anti-proliferative effect in our peptide library screen (**Fig. 2D, S2D**), highlighting the added value of our screen. For peptides that target an essential protein, and show no or little proliferation effects, we hypothesized that either these peptides were unable to outcompete essential binding partners or that they may disrupt pockets that are not critical for the essential function of the protein. Conversely, through this analysis we also identified candidate essential pockets in non-essential proteins, which we hypothesized could arise from peptides targeting analogous pockets in multiple protein homologs, inhibiting other target proteins or activating through displacement of an inhibitor. Among the 76 genetically non-essential proteins that we identified as potentially having essential peptide-binding pockets, at least 29 have paralogs, including 6 pairs that have been mapped as synthetic lethalities^25^ (**Fig. S2D**). For this reason, we chose to investigate the inhibition of TLE1-4 transcriptional co-repressor paralogs below.

### HCF1-targeting peptides reveal a therapeutic window for a genetically common essential target

As introduced earlier, we found a discrepancy between gene essentiality and sensitivity to its competitor peptides in transcriptional co-regulator HCF1. While it is genetically common essential^5,26^, sensitivity to HCF1 competitor peptides varied greatly among cell lines in our panel (**Fig. 3A-D**). HCF1 contains a Kelch domain that binds transcription factors and chromatin regulators (including proto-oncogenes such as MYC and MLL) through HCF1 binding motifs (HBM) typically containing [DENQ]HxY consensus sequences^27–29^ (**Fig. 3A-B**). Out of the 19 experimentally validated HBM peptides in the library, 11 were anti-proliferative in NALM6 (**Fig. 3D-E**). The finding that these HBM peptides, as well as three predicted HBM peptides containing a [DE]HxY consensus sequence (KANSL3, DIDO1, FOXN2) showed a greater effect in the nucleus (where HCF1 localizes) further suggests that they are likely bona fide HCF1-targeting peptides (**Fig. 3D**). While some of the HBM peptides such as KANSL3 and CREB3 are anti-proliferative in all cell lines in our panel, others including DIDO1 and THAP5, exert inhibitory effects only in a subset of lines. The sensitivity to HBM peptides did not correlate with the HCF1 gene expression or dependency. As an example, NCIH460 cells are more sensitive to peptide inhibition than CALU1 cells, despite the reverse pattern of gene dependency (**Fig. 3E, S3A**). To validate the anti-proliferative effects of HCF1-targeting peptides that we observed in the pooled dropout screen, we tested the anti-proliferative effects of individual KANSL3, DIDO1 and FOXN2 GFP-tagged peptides by flow-cytometry-based pairwise competitive growth experiments with negative control mCherry cells. Through these experiments, we saw the highest GFP/mCherry differences for peptides that had depleted the most in our pooled screen, further supporting the observations from our high-throughput screen (**Fig. 3F, S3B**).

**Figure 3.**
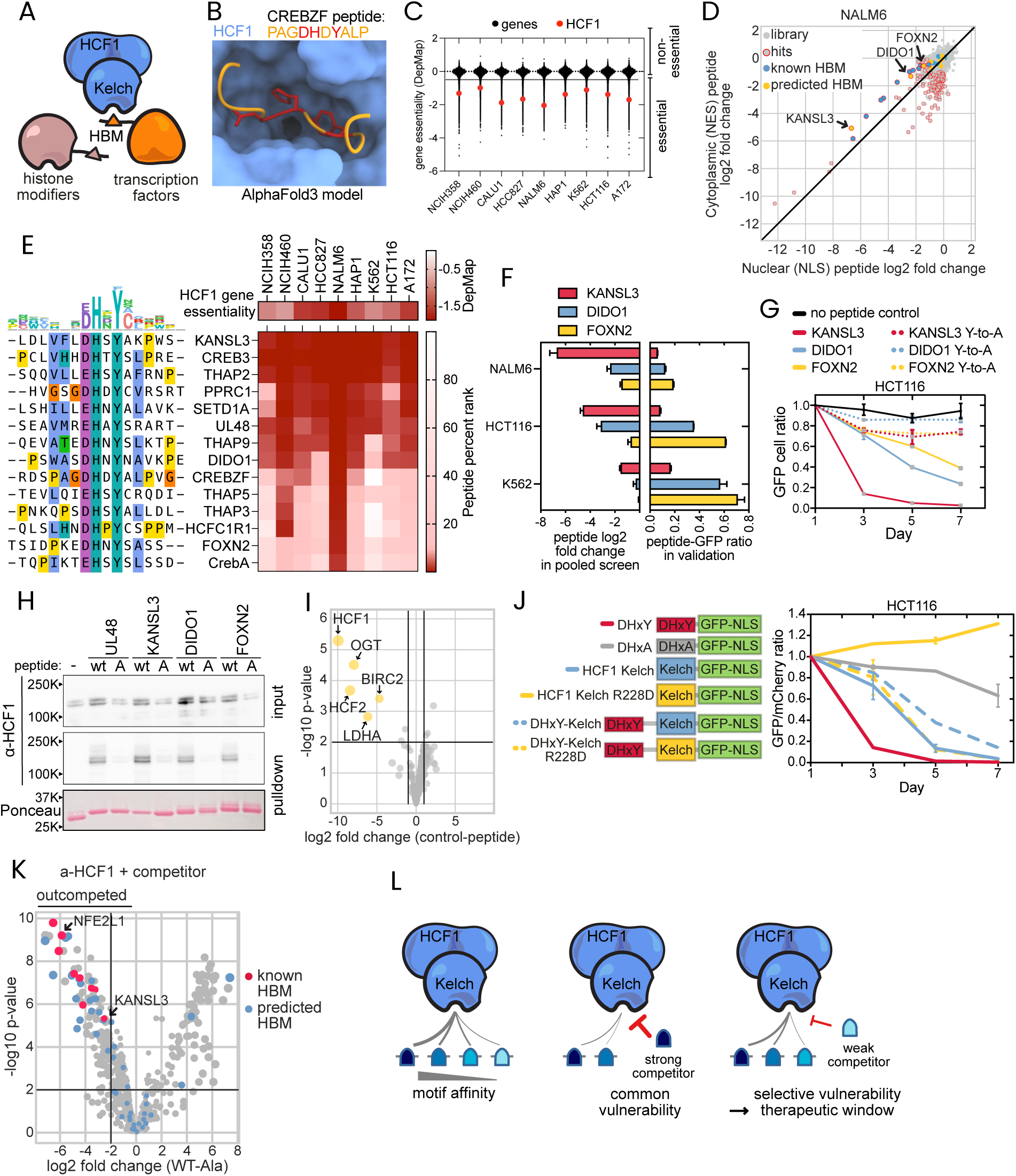
Identifying a therapeutic window by disrupting the HBM pocket of the genetically common essential Host Cell Factor-1 (HCF1). (**A**) Scheme of HCF1 binding to different proteins via the HBM-Kelch interaction. (**B**) Structural model of HCF1 Kelch domain bound to a DHxY peptide, generated with AlphaFold3. (**C**) Distribution of all gene essentiality scores (DepMap) scores in the indicated cell lines. Scores below -0.5 are considered essential. (**D**) NLS- or NES-fused peptide dropout scores measured in the proteome-wide competitive growth screen in NALM6. (**E**) Heatmap and sequence alignment of HBM peptides that were detected as an anti-proliferative peptide in at least one cell line. HCF1 gene essentiality scores are from DepMap CRISPR screens, and the red asterisks indicate essentiality (score <-0.5). (**F**) HCF1-targeting peptide dropout scores in proteome-wide screen (error bars show 95% confidence intervals of the median) and the peptide effect on proliferation in low-throughput GFP/mCherry pairwise competitive growth experiments (mean∓standard deviation). (**G**) Flow-cytometry-based pairwise competitive growth experiments showing the anti-proliferative effect of KANSL3, DIDO1 and FOXN2 peptides and the loss of peptide effect upon mutation of DHxY to DHxA. (**H**) Peptide-GFP pulldown from HCT116 lysate followed by Western blot showing the specific interaction of these peptides with HCF1. A representative example of two biological replicates is shown. (**I**) KANSL3 HBM peptide-GFP pulldown from HCT116 lysate followed by mass-spectrometry for unbiased detection of the peptide interactors. (**J**) Flow-cytometry-based pairwise competitive growth experiments showing the anti-proliferative effect of HCF1 Kelch domain overexpression and the rescue of HBM peptide toxicity by fusion to its target, the HCF1 Kelch domain. (**K**) Volcano plots showing the HCF1 interactors that are outcompeted by KANSL3 HBM competitor peptide in a HCF1 coIP from HCT116 cell lysate. (**L**) Model of HCF1-targeting competitor peptide action.

Next, we examined whether the anti-proliferative effect of these peptides arises from targeting HCF1. For this, we first introduced a single amino acid mutation in the HBM peptides (DHxY to DHxA), which effectively abolished the anti-proliferative effect of the KANSL3, DIDO1, and FOXN2 peptides (**Fig. 3G**). We then performed peptide pulldown experiments and observed that these wild-type peptides as well as a previously characterized HBM from human herpesvirus 1 protein UL48^30^ bind HCF1, whereas their tyrosine-to-alanine mutant counterparts do not (**Fig. 3H**). To test if the common anti-proliferative KANSL3 HBM peptide could have additional targets, we analyzed its interactors by proteomics and observed a highly specific interaction map with HCF1, HCF2 (an HCF1 paralog known to also bind HBMs^31^), and OGT (a binding partner of HCF1 which is necessary for its proteolytic processing and maturation) as its most enriched binders^32^ (**Fig. 3I**). Finally, as an orthogonal method to outcompete endogenous HCF1-peptide interactions, instead of overexpressing HCF1-binding peptides, we overexpressed the HCF1 Kelch domain itself. We found that the HCF1 Kelch domain overexpression had an anti-proliferative effect that is abolished with the R228D mutation in the peptide-binding pocket^33^ (**Fig. 3J**). We were able to abolish the anti-proliferative effect of the Kelch domain by fusing it to the HCF1-binding peptide, likely by intramolecular sequestering of the pocket and the peptide (**Fig. 3J, S3C**). To test if HBM peptides with different dropout scores (**Fig. 3D-E**) bind HCF1 with different strength, we used recombinant HBM peptides to pull down HCF1 from cell lysate. This revealed that KANSL3 is the strongest HCF1 interactor, followed by DIDO1 and FOXN2 (**Fig. S3D**), indicating that the peptide-HCF1 binding strength contributes to the inhibition effect size in competitive growth experiments. Together, these data suggest that these peptides inhibit cell proliferation by disrupting (to different degrees) HCF1 interactions and function.

To understand how these peptides affect HCF1 function, we next investigated the HCF1 interactome and its perturbation by the HBM competitor peptide. We found that HCF1 interacts with 441 proteins, including 43 proteins with HBMs (**Fig. S3E**), and overexpressing the KANSL3 peptide outcompeted 101 of these interactions, including 9 proteins with known HBM and 24 with potential HBMs (**Fig. 3K, S3F-H**). These outcompeted proteins are enriched in histone modifiers and transcription proteins and include four HBMs found to be essential in a base editing screen (**Fig. S3G-H**)^8^. These results suggest that KANSL3 and similar HBM competitor peptides are likely to have a profound effect on HCF1 interactions, and that the anti-proliferative effects of these peptides likely arise from the combined inhibition of a number of these interactions, rather than inhibition of any single one of them. Despite this complexity, the finding that different interactions are outcompeted to a different extent appears to create a therapeutic window where cancers more strongly relying on specific outcompeted proteins are more vulnerable to HBM competitor peptides. In line with these observations, while the cell lines in our panel exhibit considerable variation in their dependency on the proteins outcompeted by the HBM peptide, there is no considerable correlation between their dependency on any of these individual genes and their sensitivity to HBM peptides (**Fig. S3I**). Moreover, the sensitive cell lines are from diverse cancer types (**Fig. 3E**) and cannot be grouped by 50 hallmark gene sets describing cell states^34^, further suggesting a more nuanced molecular mechanism behind the sensitivity to HCF1 HBM pocket inhibition.

Finally, given the transcriptional co-regulator role of HCF1, we next performed RNAseq to further characterize the effect of HBM competitor peptides on transcription. We observed upregulation of 121 and downregulation of 359 genes (**Fig. S3J-K**). Most upregulated genes were either AP-1 or NFE2L1 targets, which aligned with our previous observation of NFE2L1 being among the top proteins outcompeted by the HBM peptide in the HCF1 interactome (**Fig. 3K**). The downregulated genes were in contrast more diverse, supporting our previous hypothesis that HBM competitor peptides likely affect a large number of processes downstream of HCF1.

Altogether our data suggest that inhibiting the Kelch domain of HCF1 can lead to a common anti-proliferative phenotype, different HBM peptides confer different levels of pocket inhibition, revealing that cancer cell lines vary considerably in their sensitivity to inhibiting this pocket (**Fig. 3E**). Given that HBM competitor peptides limit the unbound HCF1 available and that different interactors are outcompeted to different extents (**Fig. 3K**), we hypothesize that the sensitivity to HBM pocket inhibition arises from the combined partial loss of multiple interactions, where cell lines most dependent on weaker HCF1 interactors will become most sensitive to these HBM peptides (**Fig. 3L**).

### TLE-targeting WRPW peptides reveal a cancer dependency masked in genetic screens by homolog redundancy

Next, we investigated WRPW peptides binding to the WD40 domain of the transcriptional co-repressors TLE1-4^35–37^ as a candidate case of peptides targeting multiple homologous pockets (**Fig. 4A-B**). Our focus on WRPW peptides was motivated by the fact that TLE1 is an undrugged oncogene and a driver of metastasis in several cancers and that, despite TLE1-4 being non-essential genes^38–40^, several WRPW peptides were among the most anti-proliferative peptides in some cell lines in our panel (**Fig. 4C**). Moreover, the WRPW-binding pocket on TLE1-4 also binds EH1 motifs present in many undrugged cancer drivers (PAX5, FOXA1, FOXA2, TLX3, TBXT)^41–43^, further strengthening our interest into this peptide-binding pocket as a potential way to perturb multiple transcriptional programs. The effect of some WRPW peptides was highly cell line-specific, putting them among the most variable peptides in our library, and different peptides, such as the RIPPLY1/2 and the HES1 peptide showing distinct cancer cell line profiles (**Fig. 4C**). These observations led us to first ask whether the toxicity of WRPW peptides could arise from simultaneous inhibition of multiple TLE paralogs that would otherwise buffer for each other. Second, we asked if different WRPW peptides may target different sets of TLE proteins or even function through different mechanisms of action (**Fig. S4A**).

**Figure 4.**
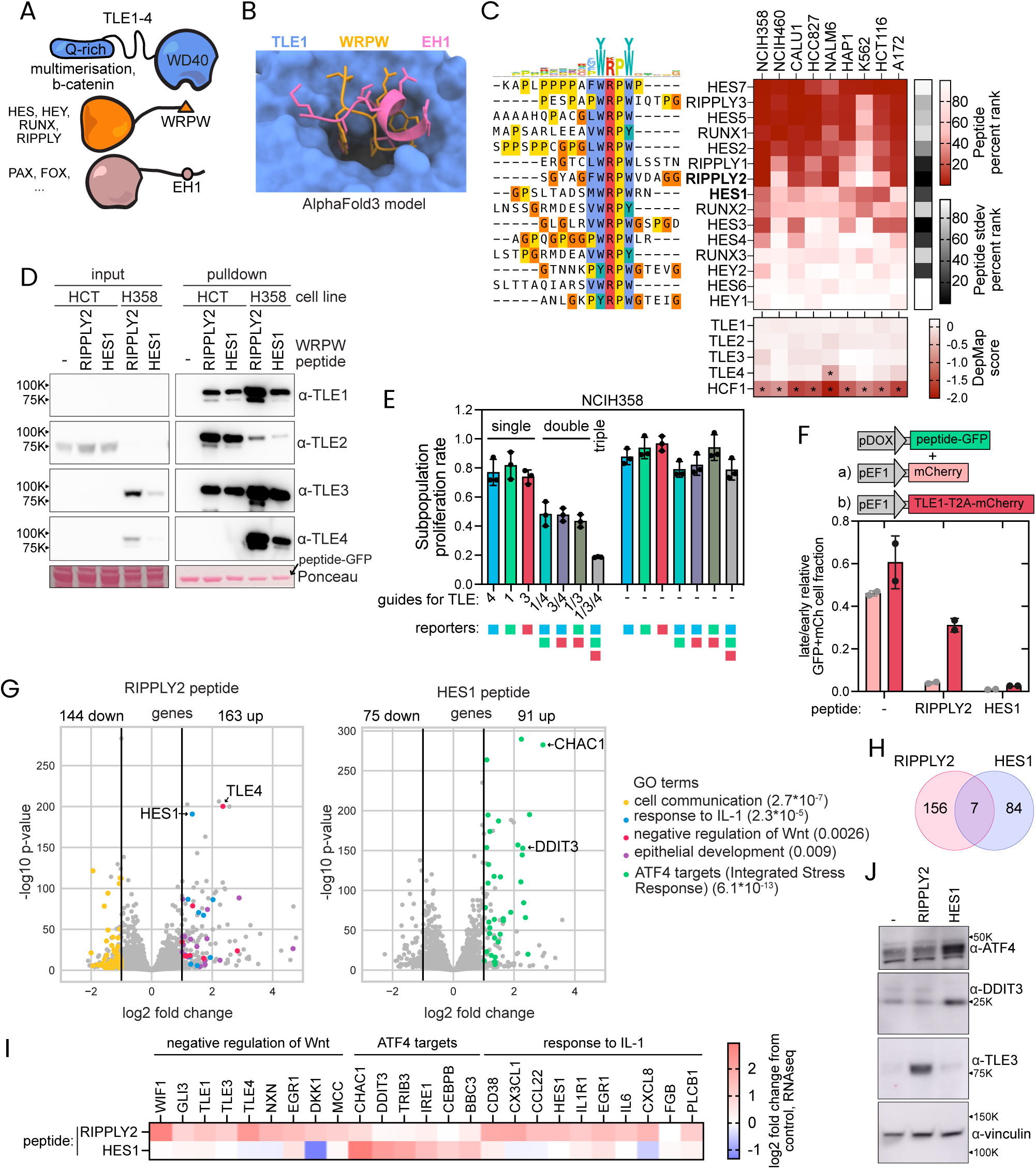
Uncovering a dependency in disrupting TLEs hidden in genetic screens by redundancy between homologs. (**A**) Scheme showing the domains in TLE1-4 proteins, where the WD40 domain binds two types of motifs - EH1 and WRPW. (**B**) Superimposed AlphaFold3 structural models of a WRPW and an EH1 peptide bound to TLE1. (**C**) Sequences and a heatmap of the dropout ranks of WRPW peptides (top) with the TLE1-4 CRISPR knockout effects on proliferation (DepMap scores, bottom). (**D**) Pulldown of RIPPLY2 and HES1 WRPW peptides from HCT116 and NCIH358 cell lysates shows binding to all four TLE1-4 proteins. A representative example of two biological replicates is shown. (**E**) The effect of single and multiplex CRISPR knockout of TLE1/3/4 in NCIH358 cells measured by the relative proliferation rates of different subpopulations co-expressing specific gRNAs and fluorescent reporters. The proliferation rates were calculated from the change in subpopulation ratios over two cell doublings. Mean±standard deviation from three biological replicates. (**F**) The impact of TLE1-T2A-mCherry overexpression on the subpopulation proliferation rate of cells expressing RIPPLY2 and HES1 WRPW GFP fusions in a flow-cytometry-based competitive growth experiment. Plot shows changes in GFP positive cell subpopulation in 5 doublings. (**G**) The impact of RIPPLY2 and HES1 WRPW peptide overexpression in NCIH358 cells investigated by RNAseq. (**H**) Numbers of unique and overlapping upregulated genes in NCIH358 cells expressing RIPPLY2 or HES1 WRPW peptides. Data is from RNAseq presented in ‘G’. (**I**) Differential effect of the WRPW peptides observed with RNAseq on ATF4 targets, genes associated with negative regulation of Wnt pathway, and genes involved in response to IL-1. (**J**) Western blot shows differential regulation of the ATF4 response and TLE3 in NCIH358 cells expressing different WRPW peptides. A representative example of two biological replicates is shown.

To test this, we first performed pulldown experiments with HES1 and RIPPLY2 WRPW peptides which differ in their cell line sensitivity profiles (**Fig. 4C**). We found that even though HCT116 and NCIH358 cells express the four TLEs to different extents, both peptides interact with and pull down all TLE1-4 (**Fig. 4D**). To test whether redundancy between TLEs had prevented genetic screens from identifying them as a cancer dependency, we next set up a multiplex CRISPR knockout experiment to study if simultaneous loss of multiple TLEs drives an enhanced anti-proliferative effect. As NCIH358 cells express TLE1, 3 and 4 more highly than TLE2 (**Fig. 4D, S4B**), we transduced the cells with vectors expressing fluorescent proteins together with sgRNAs against TLE1, 3 and 4 and monitored the proliferation of subpopulations containing those different sgRNAs over time (**Fig. 4E, S4C-D**). Consistent with previous genome-wide CRISPR screens, our data showed that loss of single TLE genes leads to minimal anti-proliferative effects^5^. In contrast, we observed a stepwise decrease in proliferation with loss of multiple TLEs, where all double knockout combinations lead to an intermediate anti-proliferative effect and TLE1/3/4 triple knockout drives the strongest anti-proliferative effect (**Fig. 4E**). Thus, this data supports our hypothesis that TLE1/3/4 are redundant in supporting proliferation, and that inhibiting them simultaneously drives a potent anti-proliferative effect.

To test if the toxicity of WRPW peptides arises from inhibition of TLE1-4, we overexpressed TLE1 and found that this partially rescues the anti-proliferative effect of the RIPPLY2 peptide (**Fig. 4F, S4E**). Although the RIPPLY2 peptide causes slight upregulation of TLE proteins (**Fig. 4D**), the observation that the RIPPLY2 peptide affected downstream TLE target proteins differently to TLE1 over-expression led us to conclude that the peptide effect is unlikely to be simply the result of target stabilization or over-expression (**Fig. S4F**). Together, our combined data suggests that the RIPPLY2 WRPW peptide inhibits multiple TLE proteins, revealing a vulnerability in a subset of cancer cell lines that has been hidden in single gene knockout or knockdown screens by paralog redundancy.

The finding that the studied cell lines differ drastically in their sensitivity to the RIPPLY2 peptide, with NCIH358, CALU1, NALM6, and A172 cells being most affected, made us ask whether we could identify common markers of sensitivity to TLE inhibition (**Fig. 4C**). We investigated the expression of hallmark gene sets in these cell lines using previously published RNAseq data^5,44^ and looked for differences between the RIPPLY2-sensitive and -insensitive cell lines. This analysis revealed that while cells with a higher oxidative phosphorylation signature tend to be more resistant to RIPPLY2 peptide, cells with a higher interferon alpha and gamma response signatures tend to be more sensitive to this TLE-binding peptide (**Fig. S4G**). In fact, TLE1-4 cofactors regulate both oxidative phosphorylation and inflammation^45^ and loss of TLE1 has been found to increase inflammatory signaling^46^, further indicating that the correlations we found may underline biologically meaningful mechanisms. Overall, our data suggests that higher inflammatory signaling may represent a mark of cells most susceptible to TLE1-4 docking inhibition.

Switching our focus onto the HES1 WRPW peptide showing distinct cell line sensitivities compared to other WRPW peptides, we found that even though this peptide interacts with TLE1-4 similarly as the RIPPLY2 peptide, the anti-proliferative effect of HES1 peptide could not be rescued by TLE1 overexpression (**Fig. 4D, F**). To better understand the different perturbations caused by these two peptides, we performed RNAseq. The RIPPLY2 peptide triggered downregulation of genes involved in cell communication and upregulation of response to interleukin-1, negative regulators of Wnt and epithelial development (**Fig. 4G**), all functions that have been associated with TLE proteins^47,48^ and further supporting a connection between TLE1-4 inhibition sensitivity and inflammatory signaling. The transcriptional response to the HES1 peptide, however, was vastly different. Despite having the WRPW motif and the ability to bind TLE1-4, the HES1 peptide did not cause upregulation of the Wnt regulators and instead drove strong upregulation of >40 ATF4 targets (**Fig. 4G-I**), indicative of activation of the Integrated Stress Response (ISR)^49^. In fact, these expression changes translate into the HES1 peptide driving an increased protein abundance of ATF4 and its pro-apoptotic target gene DDIT3, in both NCIH358 and HCT116 cells (**Fig. 4J, S4H**). Thus, while the RIPPLY2 peptide perturbations align with TLE1-4 functions, anti-proliferative effects of the HES1 peptide cannot be accounted for by TLE inhibition and are instead likely to originate from an alternative molecular mechanism of action.

### The anti-proliferative phenotype of the HES1 WRPW peptide is driven by its myristoylation and membrane localization

To identify potential additional targets of WRPW peptides that could help us elucidate the mechanism of action behind the anti-proliferative effect of the HES1 peptide, we performed a peptide pulldown from NCIH358 cell lysate followed by proteomics. Whereas the RIPPLY2 peptide primarily pulled down TLE1, TLE3, TLE4 and TLE5 with other proteins only at lower levels, the HES1 peptide pulled down additional proteins, including deoxyhypusine synthase (DHPS) and Acyl-CoA Oxidase 3 (ACOX3) at levels comparable to TLEs (**Fig. 5A, S5A**). The fact that DHPS plays a key role in maturation of eIF5a, and depletion of eIF5a leads to ATF4 activation via endoplasmic reticulum stress^50^ is consistent with the response caused by the HES1 peptide (**Fig. 4G**). We found that while both GC-7 (a small molecule inhibitor of DHPS^51^) and the HES1 peptide drove an increased ATF4 response indicated by higher ATF4 and DDIT3 protein abundance, unlike GC-7, the HES1 peptide did not decrease eIF5a hypusination (**Fig. S5B**), and overexpression of DHPS did not rescue the anti-proliferative effect of HES1 peptide (**Fig. S5C**). These results suggested that the anti-proliferative effect of the HES1 peptide was not driven by its downstream engagement of DHPS.

**Figure 5.**
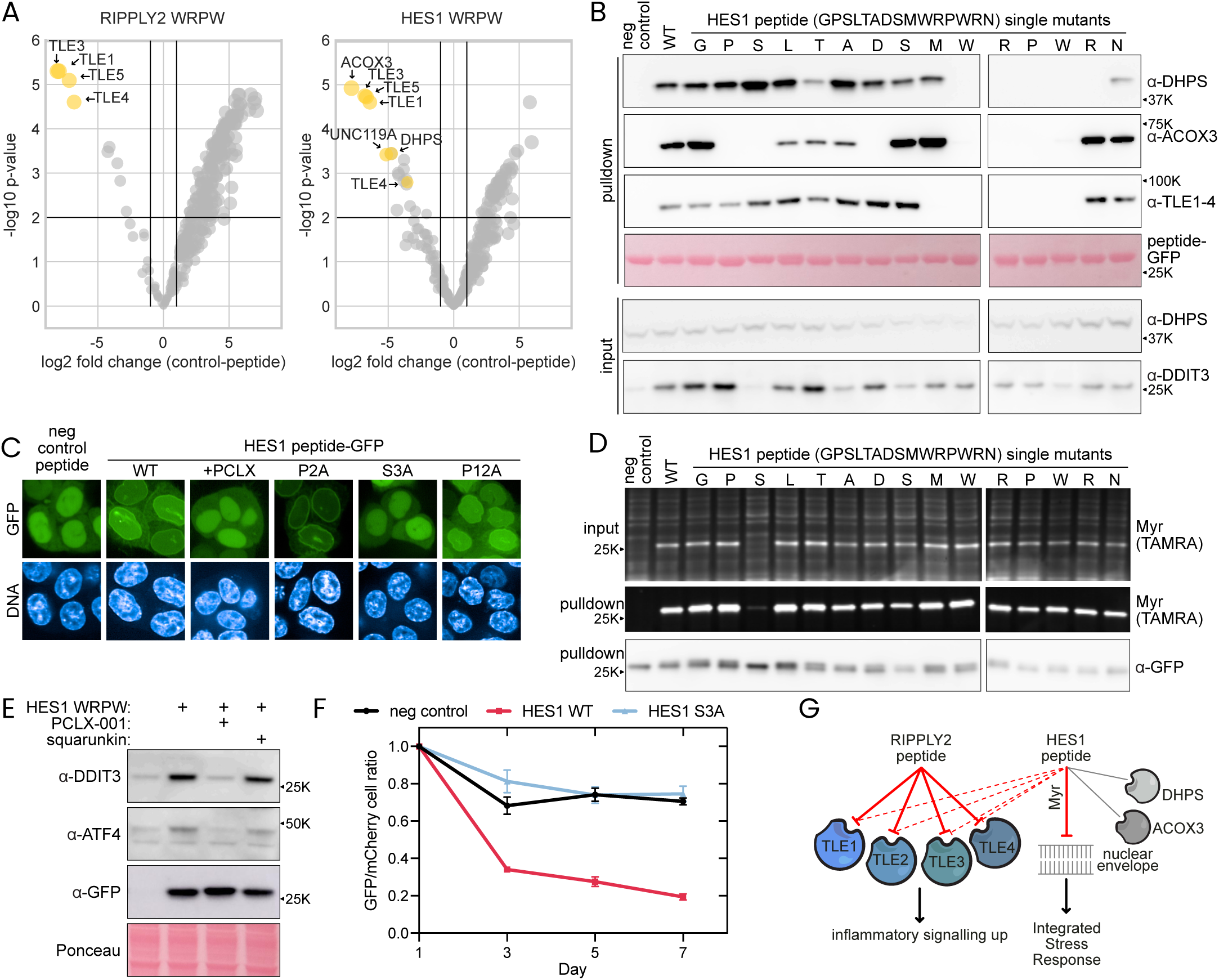
The HES1 WRPW peptide drives activation of the Integrated Stress Response by myristoylation-mediated nuclear membrane localisation. (**A**) RIPPLY2 and HES1 WRPW peptide-GFP pulldown from NCIH358 cell lysate followed by mass-spectrometry reveals distinct interactors of these peptides. (**B**) Pulldown of wild-type and single mutant HES1 peptides from HCT116 lysates showing distinct peptide binding patterns to TLE1-4, DHPS, ACOX3, and DDIT3 upregulation. (**C**) Microscopy images showing subcellular localisation of HES1-WRPW-GFP in HCT116 cells 24h after induction. PCLX-001 was used at 0.5 µM. (**D**) Myristoylation of HES1 peptide mutants monitored in HCT116 cell lysates using TAMRA-alkyne to detect azido-myristate. (**E**) Western blot showing the effect of 0.5 µM PCLX-001 and 2.5 µM squarunkin on HES1 peptide driven ATF4 response in HCT116 cells collected 24h after treatment. (**F**) Pairwise competitive growth experiment with HCT116 cells expressing either peptide-GFP or mCherry control construct. Data is mean from two independent replicates. (**G**) Model of HES1 and RIPPLY2 WRPW peptide activities.

Next, to further elucidate the molecular mechanism behind the HES1 peptide, we introduced single mutations into the peptide, aiming to obtain peptides with differences in binding and downstream phenotypes. As expected, mutations in the WRPW sequence abolished binding to TLE1-4, while most other mutations were largely tolerated (**Fig. 5B**). The experiment revealed that while binding to DHPS required a WRPWR motif, ACOX3 engagement imposed additional requirements up to eight amino acid residues N-terminal from this motif (PSxxxDxxWRPW) (**Fig. 5B, S5D**). In contrast, only the S3A mutation abolished the DDIT3 response (**Fig. 5B**). The lack of correlation between DDIT3 expression and binding to any of these associated target proteins suggested that while TLE1-4, DHPS, and ACOX3 are binders of the HES1 peptide, binding to these proteins is not behind the ATF4 response triggered by the peptide.

We found that, in contrast to the negative control that localizes to the nucleoplasm, the HES1 peptide localizes to the nuclear envelope, and this localization is abolished exclusively in the S3A mutant, in alignment with the ATF4 response (**Fig. 5B-C, Fig. S5E**). We observed UNC119A/B, which are chaperones that bind myristoylated proteins and mediate their plasma membrane localisation^52^, as HES1 peptide interactors (**Fig. 5A, S5A**). This prompted us to ask if myristoylation mediates HES1 peptide membrane localization. For this, we inhibited the N-myristoyltransferases NMT1/2 with PCLX-001^53^ and observed that this abolished the nuclear envelope localization of the HES1 peptide (**Fig. 5C, S5F-G**). Further, we observed that the S3A mutation exclusively abolished myristoylation of the HES1 peptide (**Fig. 5D**). As myristoylation, together with favorable residues flanking the modification, facilitates anchoring to membranes^54^, we propose that the myristoylated HES1 peptide binds directly to the nuclear membrane.

Consistent with this model, inhibition of N-myristoyltransferases NMT1/2 with PCLX-001 abolished the DDIT3 upregulation driven by the HES1 peptide, while the UNC119 inhibitor squarunkin had no effect (**Fig. 5E**). This suggests that the phenotype caused by the HES1 peptide is unlikely to be driven by outcompeting UNC119 interactions with its natural interactors, and instead most likely driven by the nuclear membrane accumulation of the peptide. Previous studies have found that perturbations in the nuclear envelope structure and aggregation of nuclear membrane proteins causes ISR^55,56^, providing a potential mechanism for ATF4 activation by the nuclear membrane-bound HES1 peptide. Using a pairwise competitive growth experiment, we found that the membrane-targeting activity is necessary for the anti-proliferative effect of the HES1 peptide (**Fig. 5F**). Altogether, our data indicates that the while its unexpected protein targets (eg. DHPS, ACOX3) do not appear to be necessary for the suppressive function of the HES1 peptide, its membrane-targeting activity is required for its anti-proliferative phenotype (**Fig. 5G**).

### Peptides targeting cyclin D substrate recruitment pocket reveal a leukemia-specific vulnerability

Next, with the aim to identify potential leukemia-specific peptide-binding pocket dependencies, we grouped the three leukemia cell lines (HAP1, NALM6, and K562) in our panel and compared their pocket dependencies against all other cell lines. Among the 206 peptide-binding pockets we compared, the substrate recruitment pocket of cyclin D had the greatest specificity for leukemia cell lines (**Fig. 6A**). This cyclin D pocket is known to bind to helical motifs in RB1, RBL1 and RBL2, thereby promoting substrate phosphorylation by CDK4/6 and ultimately cell cycle entry^57^ (**Fig. 6B-C**). Of these three peptides, RBL1 and RBL2 show stronger anti-proliferative effects in leukemia cell lines, whereas the RB1 peptide had weaker effect (**Fig. 6C**). The fact that in contrast to the other cell lines, NALM6 and HAP1 showed low cyclin D1 expression and dependency, leading us to investigate cyclin D expression and genetic dependency more broadly (**Fig. 6C**). DepMap data supports that solid cancers are more dependent on cyclin D1, whereas blood cancers are more dependent on cyclin D3 (**Fig. 6D**)^5^. Moreover, high cyclin D3 expression is associated with poor prognosis in acute myeloid leukemia patients (**Fig. S6A**). This led us to hypothesize that the low cyclin D1 expression and high cyclin D3 dependency could make blood cancers particularly susceptible to inhibiting the cyclin D substrate recruitment pocket.

**Figure 6.**
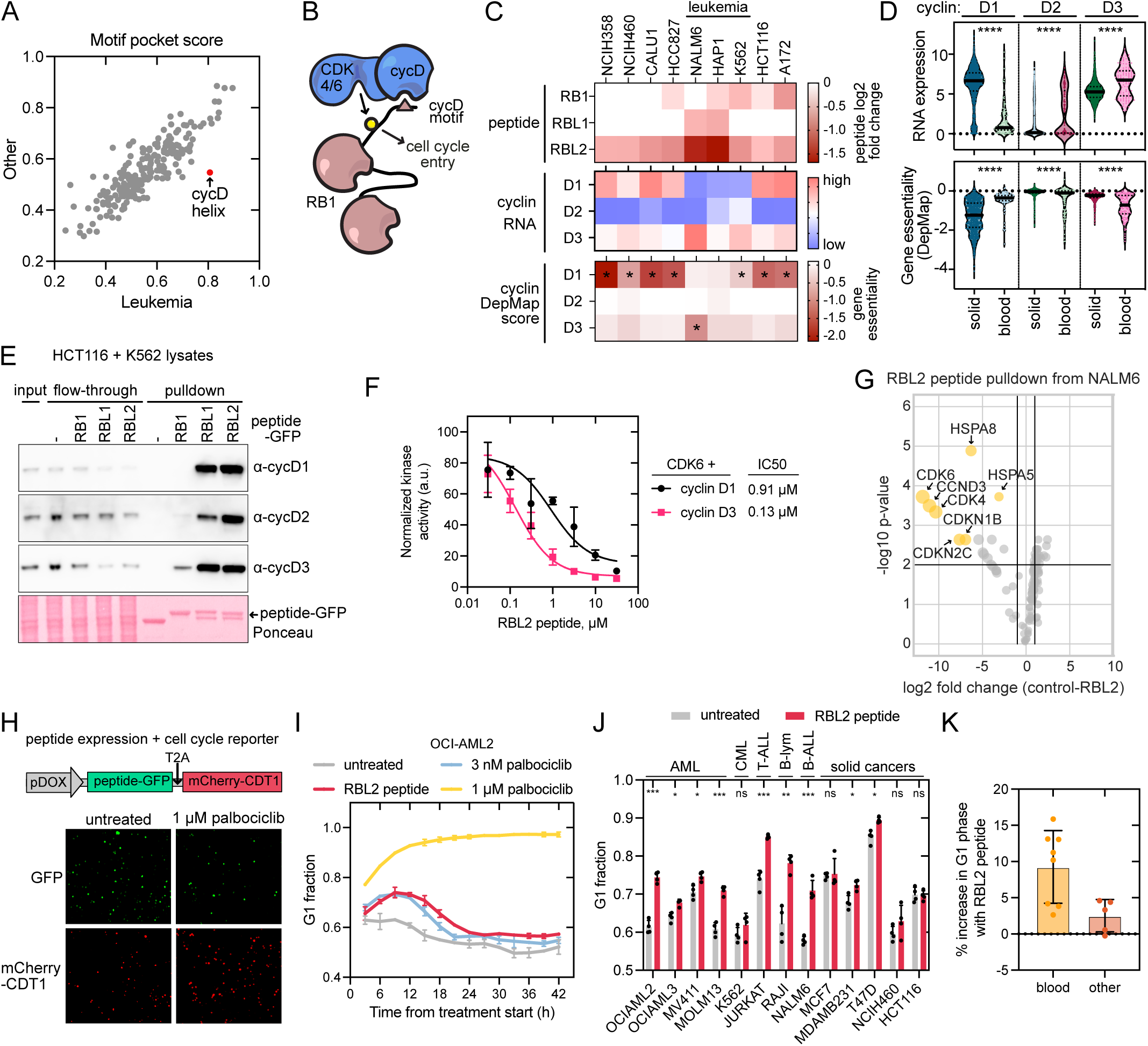
Leukemia cells exhibit increased sensitivity to cyclin D targeting peptides. (**A**) Pocket-focused analysis of motif competitor peptides reveals a leukemia-specific sensitivity for cyclin D-Rb/RBL interaction inhibition. (**B**) Scheme showing cyclin D driving CDK4/6 substrate selection by binding to helical motifs in RB1, RBL1, and RBL2. (**C**) Heatmaps showing the dropout scores of cyclin D targeting peptides in the proteome-wide screen in different cell lines (top), the expression levels of different cyclin Ds indicated by RNAseq data from DepMap (middle), and the dependencies on different cyclin Ds, indicated by CRISPR knockout scores from DepMap (bottom). (**D**) Low cyclin D1, but high D3 expression and dependency in blood cancers compared to solid cancer cell lines (DepMap data). ‘****’ indicates p-value < 0.0001 by two-sample Welch’s t-test. (**E**) Recombinant peptide-GFP pulldown using a 1:1 mixture of HCT116 and K562 cell lysates to detect binding of cyclin D targeting peptides to all three cyclin Ds. (**F**) The impact of RBL2 peptide on RbC phosphorylation rate by cyclin D1- and D3-CDK6 in vitro. Data is from three independent experiments. (**G**) RBL2 helical peptide GFP fusion pulldown followed by mass-spectrometry for unbiased identification of target proteins. (**H**) Co-expression of RBL2 peptide-GFP and mCherry-CDT1 G1-phase reporter protein enables quantification of G1-phase cells upon cyclin D-CDK4/6 inhibition by the peptides and ATP-competitive inhibitors. (**I**) Temporal dynamics of the proportion of G1-phase cells upon treatment with palbociclib measured in time-lapse microscopy using the mCherry-CDT1 reporter. (**J**) The fraction of G1-phase cells 12 hours after RBL2 peptide-GFP induction measured by mCherry-CDT1 reporter in different cancer cell lines. (**K**) RBL2-peptide driven increase in G1-phase 12h after peptide expression induction in blood and solid cancer cell lines from panel ‘J’.

To start to test this hypothesis, we first investigated which cyclin Ds these three peptides interact with. While the two anti-proliferative peptides, RBL1 and RBL2, bound all three cyclin Ds (**Fig. 6E**), the non-inhibitory RB1 peptide pulled down much lower levels of cyclin D2 and D3, and showed no detectable binding to cyclin D1 in cell lysates containing all three D-type cyclins (**Fig. 6E**). These results suggest that RB1 is a weaker cyclin D binder unlikely to be able to outcompete the endogenous cyclin D interactions, which may explain its weaker anti-proliferative phenotype. To compare inhibition of cyclin D1– and cyclin D3–CDK6 complexes, we performed *in vitro* phosphorylation assays using the C-terminal fragment of RB1 (RbC) as a substrate. Kinetic analyses demonstrated that RbC phosphorylation by CDK6 in complex with cyclin D3 was inhibited at lower RBL2 peptide concentrations than phosphorylation by the cyclin D1–CDK6 complex, with IC₅₀ values of approximately 0.1 μM and 1 μM, respectively (**Fig. 6F, S6B**). We next analyzed the RBL2 peptide interactors with proteomics to find potential additional binders. In a NALM6 lysate, the RBL2 peptide bound cyclin D3 together with CDK4/6 and CDKN1B (p27), and other potential interactors at only considerably lower signal (**Fig. 6G**), suggesting that the cyclin D complex is the main target of this peptide. Moreover, the finding that p27 is present in the RBL2 peptide pulldown suggests that the helix docking pocket is free in p27-CDK4/6-cyclin D complex and that targeting this pocket with these peptides is expected to result in inhibition of the p27-bound CDK4/6 complex, which is particularly relevant given that this complex is not targeted by palbociclib and similar CDK4/6 inhibitors^58^. Recent findings showing that a stapled peptide from RB1 is more potent than CDK4/6 inhibitors in inhibiting the p27-cyclin D-CDK4 complex are also consistent with our results^59^.

Next, as inhibiting CDK4/6 is known to lead to G1 arrest, we investigated the impact that the cyclin D-targeting peptides have on cell cycle progression. By propidium iodide (PI) DNA staining and flow cytometry analysis in the leukemia cell line NALM6, we found that the RBL1/2 peptides cause an increase in G1-phase cell fraction, albeit to a lesser extent than the CDK4/6 inhibitor palbociclib (**Fig. S6C**). To understand the temporal profile of the response to cyclin D-targeting peptides, we set up live cell time-lapse imaging experiments using mCherry-CDT1, the G1-phase reporter from the FUCCI system^60^ (**Fig. 6H, S6D**). These experiments revealed that the RBL2 peptide causes a transient increase in G1-phase cells, peaking at around 9-12 hours after inducing peptide expression (**Fig. 6I**). This temporal profile is similar to the one seen with low concentrations (3 nM) of palbociclib, while at higher concentrations palbociclib causes a more stable and complete G1-phase arrest (**Fig. 6I**).

When we expanded the number of cancer cell lines, we observed that most tested blood cancer cell lines were sensitive to expression of cyclin D targeting peptides and showed an increased G1 fraction post peptide induction (**Fig. 6J**). In contrast the peptide had no or little effect on most other cell lines, including breast cancer lines (**Fig. 6J-K**). Thus, we conclude that a large fraction of blood cancers may be specifically sensitive to inhibiting the cyclin D docking pocket compared to solid cancers, possibly arising from their stronger dependency on cyclin D3, which is more sensitive to peptide inhibition compared to cyclin D1.

### Cyclin D-targeting peptides enhance the effects of ATP-competitive CDK4/6 inhibitors

Given that cyclin-substrate interactions often drive substrate phosphorylation and that this docking pocket appears to be targetable independently to the kinase active site, we next tested if treating cancer cells with the RBL2 peptide would increase their sensitivity to palbociclib. Using time-lapse imaging with the mCherry-CDT1 G1-phase reporter, we measured the IC50 of palbociclib in different cancer cell lines expressing either RBL2 peptide fused to EGFP or a no peptide EGFP control construct. We observed that the RBL2 peptide expression decreased the IC50 of palbociclib 20-fold and 118-fold in breast cancer lines MDA-MB-231 and T47D, respectively (**Fig. 7A**), even when we had observed that the peptide alone had only a small effect in these cells (**Fig. 6J**). The RBL2 peptide also decreased the IC50 of palbociclib in leukemia cells, albeit to a smaller 2- to 6-fold extent (**Fig. 7A**). Given how much cancer cell lines differ in the extent of palbociclib IC50 improvement with the peptide, our results suggest that active site and docking pocket co-targeting could be leveraged to improve treatment specificity. More specifically, the finding that RBL2 peptide improves the IC50 of palbociclib over 20-fold in breast cancer line MDA-MB-231 while increasing it only up to 6-fold in blood cancer cells suggests that the co-targeting treatment may enhance potency against breast cancer while potentially relieving the hematopoietic adverse effect of palbociclib that limit current treatment^61^. We also found that the RBL2 peptide expression decreases the IC50 of atirmociclib, a CDK4 specific inhibitor that has shown improved treatment specificity in preclinical experiments^62^ (**Fig. 7B**). While the IC50 of atirmociclib was 3.7- and 4.4-fold lower in MCF7 breast cancer cells compared to MV-4-11 and NALM6 leukemia models, respectively, the combined treatment of RBL2 peptide and atirmociclib showed over 7.6-fold higher sensitivity in MCF7 compared to the blood cancer cells (**Fig. 7B, S7A,B**). Thus, our experiments show that dual targeting of the cyclin D-CDK4/6 complex with ATP-competitive drugs and docking pocket-targeting inhibitors could improve treatment efficacy and specificity.

**Figure 7.**
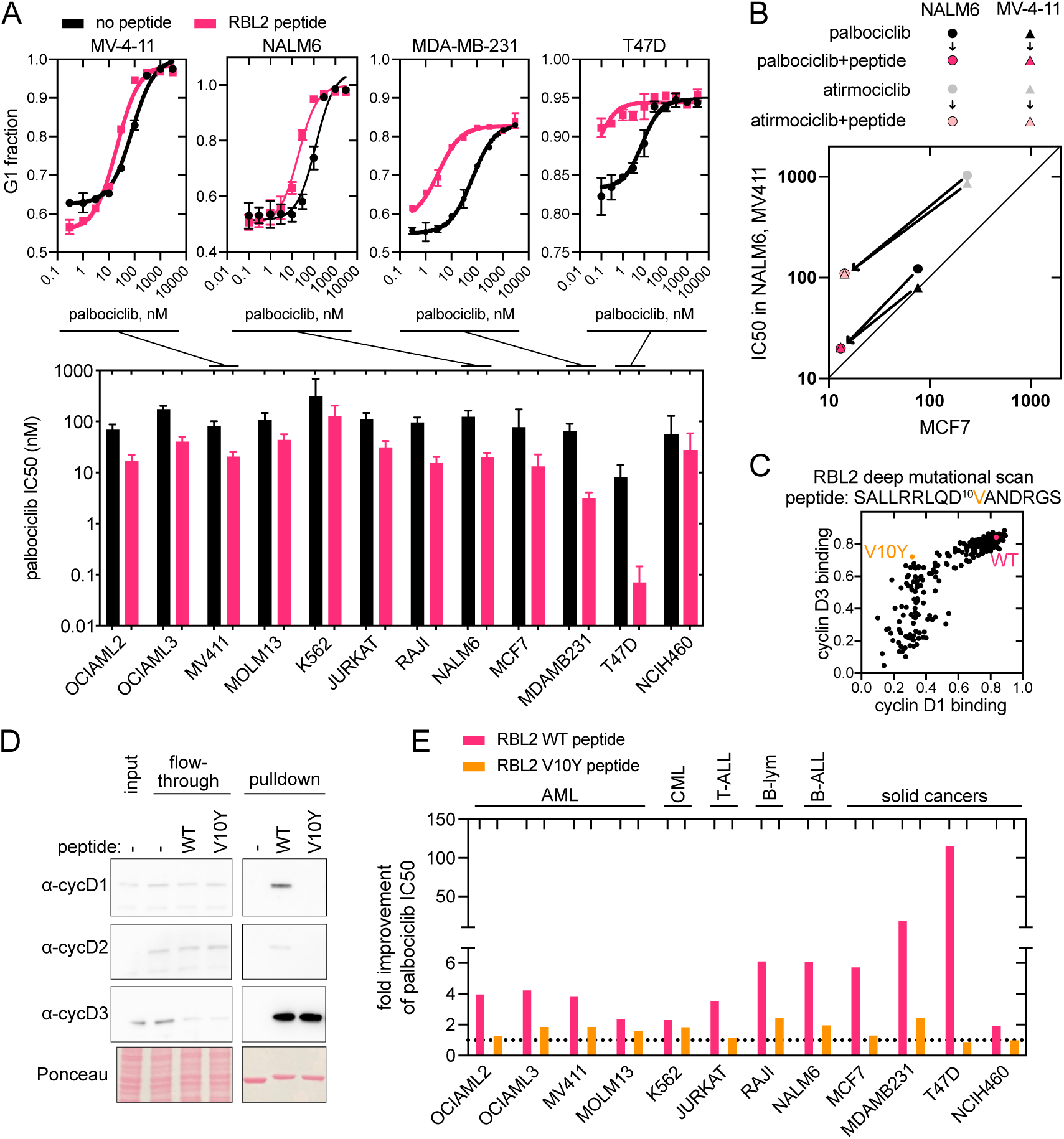
Cyclin D targeting peptides improve the effect of ATP-competitive CDK4/6 inhibitors. (**A**) Top: dose-response curves showing the fraction of G1-phase cells 24h after start of treatment with RBL2 cyclin D targeting peptide and palbociclib. Bottom: palbociclib IC50 values in different cancer cell lines measured by the fraction of G1-phase cells 24h after treatment in the presence or absence of the RBL2 competitor peptide. Error bars show 95% confidence intervals. (**B**) IC50 of palbociclib and atirmociclib in the presence or absence of the RBL2 peptide in MCF7 cells compared to NALM6 and MV411 cells. (**C**) Deep mutational scanning of RBL2 peptide combined with cyclin D1 and D3 binding measurements in SIMBA reveal mutations that lose cyclin D1 binding, but retain D3 binding (V10Y). (**D**) Recombinant peptide-GFP pulldown using a 1:1 mixture of HCT116 and K562 cell lysates to detect binding of cyclin D targeting peptides to all three cyclin Ds. (**E**) The effect of wild-type or V10Y mutant RBL2 peptide on palbociclib IC50 in different cancer cell lines measured by the G1-phase cell fraction 24h after treatment.

While the wild-type RBL1/2 peptides interacted with all three cyclin Ds (**Fig. 6E**), different cancers rely on different cyclin D genes (**Fig. 6D**), which led us to examine if the cyclin D docking pocket could be targeted in a cyclin-specific manner. To this end, we investigated a deep mutational scanning (DMS) dataset of the RBL2 peptide^59^ to identify peptides with preferential binding to either cyclin D1 or D3. This analysis revealed that binding to cyclin D3 is more permissive of mutations in the RBL2 peptide than binding to cyclin D1, effectively resulting in peptide variants such as RBL2^V10Y^, which retain binding to cyclin D3, while significantly losing cyclin D1 binding (**Fig. 7C**). To validate these results in a human cell lysate setting, we performed peptide pulldown experiments from cell lysates, where we observed that RBL2^V10Y^ peptide specifically pulls down cyclin D3, and not cyclins D1 or D2 (**Fig. 7D**). These results confirm that targeting this pocket has the potential to introduce cyclin-specificity into CDK4/6 inhibitory treatments. Next, we tested the impact of the cyclin D3-specific RBL2^V10Y^ peptide on the IC50 of palbociclib and found that RBL2^V10Y^ peptide decreased the IC50 in all leukemia cell lines, albeit to lesser extent than the RBL2^WT^ peptide (**Fig. 7E**). The RBL2^V10Y^ peptide effect varied more than the wild-type peptide effect among different leukemia lines, being greatest in RAJI and weakest in JURKAT cells (**Fig. 7E**). Moreover, while the palbociclib sensitivity of some solid cancer cell lines (such as T47D and NCIH460) was not affected by the RBL2^V10Y^ peptide, as expected given their genetic dependency on cyclin D1 and not cyclin D3 (**Fig. S7C,D**), other breast cancer cell lines (such as MDA-MB-231 and MCF7) were affected by the RBL2^V10Y^ peptide, albeit to much lesser extent than by the wild-type peptide (**Fig. 7E**). Overall, we observed that the RBL1^V10Y^ peptide was relatively more potent in improving palbociclib IC50 in blood cancer cells compared to the tested breast cancer cells (**Fig. S7E**), suggesting that cyclin-specific targeting of cyclin D-CDK4/6 complexes could also be further explored as an approach to improve treatment specificity.

## Discussion

In this study, we investigate the phenotype of inhibiting specific protein-protein interaction pockets using peptides that outcompete endogenous interactions. Similar to genes that have been categorized as common essential, specific, or non-essential based on data from genetic screens^5,6,63^, we find that inhibiting some peptide-binding pockets is anti-proliferative across all cell lines, while inhibition of others can either have cancer-type, cell-line-specific anti-proliferative effects or no significant effect in any cell line (**Fig. 2A**).

By comparing our screening results to previous genetic screens, we find that often the pocket dependency does not correlate with target gene dependency, as we find both non-essential pockets in essential proteins and highly anti-proliferative peptides targeting non-essential proteins. This difference from CRISPR gene knockout studies may arise through multiple mechanisms. First, the competitor peptides tend to only inhibit selected functions and interactions of any given target. For example, while the HCF1-targeting competitor peptides outcompete at least 32 HCF1 interactors, other interactions, such as the HCF1-OGT interaction, remain intact (**Fig. 3I-K**). Second, the competitor peptides also tend to lead to partial inhibition of a given pocket, instead of a complete inhibition. In fact, such dose effects have also been observed in gene dependency studies, where 50% of genes identified as common essential by CRISPR were found to be selective dependencies by RNAi which, unlike CRISPR, drives a partial loss of expression^64^. As most pockets interact with peptides from multiple proteins that are affected to a different extent by the competitor peptides, this effectively leads to pocket inhibition sensitivity profiles that differ from gene dependency profiles and cell lines having different sensitivity to inhibitors for a pocket in a common essential protein. As an example, we identified the peptide-binding pocket on the Kelch domain of HCF1 (a common essential gene) as an actionable pocket after seeing how cell lines varied greatly in their sensitivity to HCF1-targeting peptides. The fact that the oncogenic potential of MYC is dependent on its ability to interact with HCF1 via the Kelch domain^65^ further emphasizes how these results may reveal new ways to target oncogenic proteins. As many intrinsically disordered proteins that lack druggable pockets rely on motifs for protein-protein interactions, inhibiting peptide-binding pockets on their binding partners may enable the (indirect) inhibition of disordered oncogenes.

Another key difference of our peptide technology from more traditional gene knockout screens is that peptides can, like small molecule drugs, inhibit multiple targets simultaneously. Such multi-targeting can occur for targets that share homologous pockets, such as TLE1-4 targeted by the RIPPLY2 peptide (**Fig. 4**), or arise from the ability of a peptide to bind and inhibit multiple phylogenetically unrelated proteins, such as TLEs, DHPS, and ACOX3, targeted by the HES1 peptide (**Fig. 5**). Both of these types of multi-target specificity have been observed and shown to be essential for the efficacy of specific drugs. For example the potency of imatinib depends on inhibition of multiple kinases^66^ and several drug candidates have been found to exert their anticancer effect by “off-target” inhibition^67,68^. Given that the majority of human genes have paralogs and paralogous genes are less likely to be essential^69,70^, gene redundancy is likely to be common and could lead to potentially valuable targets being untapped from single gene knockout screens. Consistent with this idea, in recent years, CRISPR screens targeting pairs of paralogs have identified additional cancer vulnerabilities^69,71,72^. Moreover, a fraction of peptides in our library likely target other proteins in addition to their characterized binding partner. One such peptide is the HES1 peptide which contains at least 5 different functionalities including overlapping, but not identical binding motifs for its characterized TLE1-4 targets, as well as DHPS and ACOX3, a myristoylation motif and residues that appear to promote interactions with the nuclear membrane (**Fig. 5**). While this illustrates the complexity of competitor peptide screens, it also highlights the potential of encoding different functions into short peptides.

Many drugs hit catalytic sites of enzymes that are often conserved among many family members (which hinders target selectivity) and readily evolve resistance mutations. As an orthogonal approach, targeting peptide-binding pockets would expand the druggable proteome space and provide opportunities for enhanced selectivity and alternative pockets to overcome resistance. We found that leukemia cells are more sensitive to peptides inhibiting cyclin D interactions (**Fig. 6**), providing additional means to inhibit the CDK4/6 complex, a major therapeutic target in several cancers^73^. Moreover, we showed that peptides targeting cyclin D, a substrate adaptor subunit for CDK4/6, greatly enhance the sensitivity to ATP-competitive CDK4/6 inhibitors (**Fig. 7**). Simultaneous drugging of multiple pockets such as catalytic and allosteric sites, has already proven successful in other kinases with benefits arising from increased target specificity by combining the selectivity of multiple pockets, additional decreased off-target toxicity due to using lower drug doses, and lower risk of resistance^74,75^.

Our study provides a resource of anti-proliferative phenotypic data for the inhibition of different peptide-binding pockets and may serve as a starting point for the design of small molecule drugs. Recent advances in mRNA and DNA cellular delivery enabling the genetic expression of therapeutic proteins open up the possibility to use peptides even directly as therapeutics^76–78^. With less than 10% of motif peptides characterized to date^14,15^, recent progress in high-throughput motif identification^79^ will likely increase the pool of well-characterized peptides available for screening of future therapeutic possibilities. Moreover, methods like DMS enable the optimization of binding strength and specificity of peptides, facilitating the design of peptides targeting distinct sets of homologous targets^80,81^. Together with phenotypic screening of these peptides in cells, these approaches could greatly accelerate our knowledge around the therapeutic potential of targeting different pockets and the importance of target specificity. In doing so peptide-mediated interactions will likely be leveraged to target and treat pathologies, infectious diseases (where viruses are known to hijack and rely on human peptide-binding pockets) and cancer^82–84^.

## Data and Code Availability

The raw data generated in this study is available in the supplementary material.

The python code used for data processing in this study is publicly available and has been deposited in Github at https://github.com/Creixell-lab/motif-pocket-dependencies, under Apache 2.0 license.

## Acknowledgements

We would like to thank the Proteomics, Genomics, Research Instrumentation and Cell Services, Flow Cytometry, Microscopy and Bioinformatics core facilities at Cancer Research UK Cambridge Institute for their critical support and contribution to this project. We also thank Peter Pryciak, Rimma Belotserkovskaya, Jordan Wilson, Soleilmane Omarjee, Ian Cannell, Matthew Ford, Louise O’Brien, Konstantinos Tzelepis, and all members of the Creixell lab for insightful discussions and for sharing reagents. We thank Eli-Eelika Esvald for preparing the purified enzymes and all members of the Koivomagi lab for helpful discussions. This research was funded by UKRI (EP/X042065/1 to M.O.), Blood Cancer UK (Innovative Pilot Grant 25016 to M.O., P.C.), the Intramural Research Program of the National Institutes of Health (NIH Grant ZIA BC 012133 to M.K.), Cancer Research UK Senior Cancer Research Fellowship (C68484/A28159 to N.E.D.), core support from Cancer Research UK (CRUK C9545/A29580 to P.C.), Children’s Brain Tumour Centre of Excellence (C9685/A26398 to P.C.), and The Brain Tumour Charity (GN-000758 to P.C.). The contributions of the NIH author(s) are considered Works of the United States Government. The findings and conclusions presented in this paper are those of the author(s) and do not necessarily reflect the views of the NIH or the U.S. Department of Health and Human Services.

## Author contributions

Conceptualization, M.O., N.E.D and P.C.; Methodology, M.O., M.K.; Software, M.O.; Investigation, M.O., L.K., D.B., M.L., M.K.; Writing—original draft, M.O.; Writing—review & editing, M.O., N.E.D. and P.C. with contributions from all authors; Funding acquisition, M.O., M.K., N.E.D. and P.C.

## Methods

### Cell lines and culturing

HCT116, NCIH358, NCIH460, CALU1, HCC827, A172 were obtained from ATCC. HAP1 C859 was obtained from Horizon Dynamics. OCIAML2, OCIAML3, MV-4-11, MOLM13 were kind gifs from Konstantinos Tzelepis lab, JURKAT and Raji cells were kind gifs from Maike de la Roche lab, MCF7, MDA-MB-231 and T47D were kind gifts from Jason Carroll lab. HCT116 was cultured in McCoy 5A medium (36600021, Gibco), NCIH358, NCIH460, CALU1, HCC827, NALM6, MV-4-11, MOLM13, JURKAT, Raji, MCF7, MDA-MB-231, and T47D in ATCC-modified RPMI 1640 medium (11504566, Gibco), A-172 and HEK293T in DMEM with pyruvate (41966029, Gibco), K562 in IMDM (12440053, Gibco), and OCIAML2 and OCIAML3 in MEM ɑ with nucleosides (12571063, Gibco). All media were supplemented with 10% fetal bovine serum (A5209402, Gibco), 100 U/ml penicillin, 0.1 mg/ml streptomycin (15140122, Gibco). All cell lines were cultured in a 37 °C incubator with 5% CO_2_. All cell lines were tested negative for Mycoplasma by qPCR at the Cancer Research UK Cambridge Institute Research Instrumentation and Cell Services core facility.

Doxycycline hyclate (D9891-1G, Sigma-Aldrich) was dissolved at 1 mg/ml in DMSO. Nutlin-3a (S8059, Selleck) was dissolved at 5 mM in DMSO. GC7 (HY-108314A, MedChemExpress) was dissolved at 10 mM in H_2_O. PCLX-001 (T63788, Cambridge Bioscience) was dissolved at 1 mM in DMSO. Squarunkin A (HY-127002, Cambridge Bioscience) was dissolved at 5 mM in DMSO. Palbociclib (S1116, Selleck) was dissolved at 10 mM in DMSO. Atirmociclib (PF-07220060, M45136, Clinisciences Limited) was dissolved at 5 mM in DMSO.

### Generation of overexpression cell lines

The competitor peptide and EGFP coding sequences were cloned into doxycycline-inducible lentiviral vector pCW57.1 (Addgene #41393) using NEB Stable Competent *E. coli* cells, followed by plasmid DNA extraction with QIAfilter plasmid kits (Qiagen). The virus was produced as follows: 7 × 10^5^ HEK293T/17 cells (ATCC: CRL-11268) were seeded to 6-well plate and in 24 h were transfected using lipofectamine 3000 (Thermo Fisher Scientific) with psPax2 (Addgene #12260) and pMD2.G (Addgene #12259) as packaging plasmids. 6 h after the transfection, the medium was replaced with 3 ml fresh DMEM. For adherent cell lines, 2-3 × 10^5^ cells were seeded to a 6-well plate 24 h before transduction. The medium containing the virus was collected 2 days after transfection and together with 8 µg/ml polybrene (TR-1003-G, Sigma-Aldrich) was used to transduce the recipient cells. For suspension cell lines, the viral supernatant was mixed with 2 million cells and 8 µg/ml polybrene, followed by centrifugation for 2h at 1000g 30 °C. In two days, the transduced cells were split and supplemented with 1 µg/ml puromycin to eliminate untransduced cells. Prior to seeding for experiments, the transduced cells were split at least 3 times and cultured in the presence of 1 µg/ml puromycin (A1113803, Gibco) to allow selection of the transduced cells. To construct NCIH358 Cas9 cell line, lentiCas9-Blast (Addgene #52962) was transduced similarly as described above, with 5 µg/ml blasticidin (SBR00022, Sigma-Aldrich) used for selection.

### Immunoblotting

Cells were collected by trypsinization, washed with PBS and snap frozen. For immunoblotting, the cell pellets were resuspended in a buffer containing 10 mM Tris-HCl pH 7.4, 150 mM NaCl, 1% Triton X100, phosSTOP phosphatase inhibitor cocktail (Roche) and cOmplete EDTA-free protease inhibitor cocktail (Merck) and pulled through a 26G needle. The lysate was cleared by centrifugation and the total protein concentration was measured using Pierce™ Bradford Plus Protein Assay Reagent. For immunoblotting, 20-40 µg of cell lysate was resolved using SDS-PAGE and the proteins were transferred to nitrocellulose membrane using iBlot 3 (Thermo Fisher Scientific). After transfer, the membrane was blocked using 5% fat-free milk solution in TBS-T, followed by overnight incubation at 4 °C with the primary antibody solutions. The following primary antibodies were used: ab13970 chicken polyclonal to GFP at 1:5000, p21 (12D1) Rabbit mAb (#2947, Cell Signaling Technologies) at 1:1000, p53 (1C12) Mouse mAb (#2524, Cell Signaling Technology) at 1:500, Hcfc1 Antibody (Amino-terminal Antigen) (#69690, Cell Signaling Technology) at 1:1000, TLE1 (F-4) (sc-137098, Santa Cruz Biotechnologies) at 1:500, TLE2 (D-10) (sc-374226, Santa Cruz Biotechnologies) at 1:500, TLE3 (D-10) (sc-514798, Santa Cruz Biotechnologies) at 1:500, TLE4 (E-10) (sc-365406, Santa Cruz Biotechnologies) at 1:500, TLE1/2/3/4 (#4681, Cell Signaling Technologies) at 1:500, ATF4 (PA5-27576, Thermo Fisher Scientific) at 1:2000, DDIT3 (R-20) (sc-793, Santa Cruz Biotechnologies) at 1:500, Vinculin (#13901, Cell Signaling Technology) at 1:1000, DHPS (A-10) (sc-365077, Santa Cruz Biotechnologies) at 1:500, ACOX3 (17360-1-AP, Proteintech) at 1:500, cyclin D1 (sc-20044, Santa Cruz Biotechnologies) at 1:500, cyclin D2 (D52F9) (#3741, Cell Signaling Technology) at 1:1000, cyclin D3 (DCS22) (#2936, Cell Signaling Technology) at 1:1000, eIF5A (D8L8Q) (#20765, Cell Signaling Technology) at 1:1000, hypusine (ABS1064-I, EMD Millipore) at 1:1000. Then, the membrane was washed 5 times with TBS-T, incubated with secondary antibody solutions for 1 hour, and washed again 5 times with TBS-T. HRP-conjugated anti-mouse IgG (#7076, Cell Signaling Technologies) at 1:3000, HRP-conjugated anti-rabbit IgG (#7074, Cell Signaling Technologies) at 1:10000, and HRP-conjugated goat anti-chicken IgY (ab97135, Abcam) at 1:10000 were used as secondary antibodies. The antibodies were detected using SuperSignal™ West Pico PLUS Pico Chemiluminescent substrate (Thermo Fisher Scientific) and Amersham ImageQuant.

### Flow-cytometry-based competitive growth experiments

For pairwise competitive growth experiments, lentiviral constructs for doxycycline-inducible peptide-GFP fusions were transfected with a control construct expressing mCherry at equimolar ratio into HEK293T/17 cells as described above. The viral supernatant was used to transduce the recipient cell lines at multiplicity of infection (MOI) ∼0.2-0.5 so that the majority of cells would receive either the GFP or the mCherry construct. The transduced cells were selected with 1 µg/ml puromycin for at least 1 week, followed by addition of doxycycline to start peptide expression and the pairwise competitive growth. The first time point was 24h after treatment start, followed by three sequential ⅙ split and cell collections at 80% confluency. During the experiment, the doxycycline-containing medium was replaced every 48h. In Fig. 3F and S3B different doxycycline concentrations were used for different cell lines to obtain comparable peptide-GFP expression levels: 0.05 µg/ml for HCT116, 0.5 µg/ml for K562, and 2 µg/ml for NALM6.

For multiplex TLE1/3/4 CRISPR knockout competitive growth experiments, sgRNA-expressing constructs (LentiGuide-puro-NLS-GFP (Addgene #185473), LentiGuide-puro-NLS-mCherry (Addgene #185474), LentiGuide-puro-NLS-BFP, TLE1 gRNA: CACGATGCAGAGCACCACAG; TLE3 gRNA: GCCGGGATTTAAATTCACGG; TLE4 gRNA: AAGCCCGGCACTGCTACCGA)^86^ were transduced to NCIH358 Cas9 cell line at high MOI to obtain cells with expressing multiple gRNAs and fluorescent reporter proteins. The first time point was collected 3 days after transductions, followed by cell collections and 1/4 splits when cells reach 80% confluency.

For all experiments, the collected cells were washed with PBS and analyzed on BD LSR Fortessa flow cytometer to quantify cell populations expressing different fluorescent reporters.

### Design and cloning of proteome-wide motif competitor peptide library

The proteome-wide motif competitor peptide library contained experimentally validated peptides and predicted motif peptides. The experimentally validated peptides were extracted from manually curated databases (MoMap, ELM^15^) or from PDB structures containing disordered peptides. The motif predictions were performed with the SliMPrints^87^ algorithm to identify sequences with higher local conservation in intrinsically disordered regions. The predicted motifs were filtered with two cut-offs: a SLiMPrints significance score of 0.005 was used for motifs in essential proteins (based on DepMap mean gene effect < -0.5) or for motifs that overlap disease relevant SNVs, while a significance score of 0.0005 cut-off was used for other motifs.

This initial set of motif peptides was subjected to multi-level filtering: 1) keeping only human peptides or natural peptides shown to bind human proteins; 2) keeping peptides with length 3-15 amino acids; 3) dropping peptides known to function as terminal peptides (e.g. PDZ motifs); 4) removing N-glycosylation and mannosylation peptides; 5) dropping extracellular, lumenal, transmembrane, mitochondrial peptides, and peptides from folded domains, based on PepTools annotations^88^. Overlapping peptides that fit into a single 15mer were merged. Peptides shorter than 15 residues were made into 15mers to include flanking sequences from the parent protein. The library was supplemented with a negative control peptide set containing 100 peptides with randomly chosen amino acids in each position and 101 peptides, where key known binding determinants were mutated to alanine. The alanine mutant control set consisted of PCNA_PIP, ANK_Tankyrase, SWIB_MDM2, and Cyclin_Hydrophobic_Patch_RxL peptides.

The peptides were expressed as two motif repeats, followed by a TRTGSTGS linker, EGFP, and an SV40 nuclear localisation (PKKKRKV) or a CRM1-dependent nuclear export signal (LALKLAGLDIGS). These expression cassettes were cloned into pLV-EF1a-IRES-Puro (Addgene #85132) based lentiviral vector with pEF1a promoter. The first motif repeat was encoded using most likely codons, while the second repeat was encoded by randomly chosen codons that differed from the first repeat. When possible, the GC content of the peptide-encoding sequences was between 40-60% and homopolymers longer than four nucleotides were not allowed. The oligonucleotides were ordered as a pooled library from GenScript. The oligonucleotide library was PCR-amplified, digested with BamHI-HF and MluI-HF (NEB), ligated into pLV-EF1a-EGFP-NLS/NES, and transformed into NEB Stable Competent *E. coli* cells, obtaining 135 times number of colonies over variants in the library.

### Pooled competitor peptide library screening

HEK293T/17 cells were transfected with the pooled lentiviral expression plasmid libraries (pLV-EF1a-peptide-EGFP-NLS/NES) and psPax2 and pMD2.G as packaging plasmids using lipofectamine 3000. 6 h after the transfection, the medium was replaced with fresh DMEM. The viral supernatant was collected by filtering through a 45 µm nitrocellulose filter first at 48 h after transfection, when fresh DMEM was added to the HEK293T/17 cells for second viral supernatant collection in 24 h. The virus was used to transduce 20 million recipient cells (∼1400x over the number of variants in the library) at MOI 0.2-0.5. For adherent cell lines, the viral supernatant was added to the recipient cells together with 8 µg/ml polybrene. For suspension cell lines, the viral supernatant was mixed with the recipient cells and 8 µg/ml polybrene, followed by centrifugation for 2 h at 1000 g 30 °C. In two days, the transduced cells were trypsinized to collect half of the cells for the t0 time point and to seed the other half of the cells with 1 µg/ml puromycin for the competitive growth experiment. A small fraction of cells was analysed by flow cytometry for EGFP expression to confirm MOI. Cells were cultured for ∼8 doublings for the competitive growth experiment (in medium containing 1 µg/ml puromycin), splitting cells ⅙ for two times when cells reached 90% confluency. The cell culture medium was replaced every three days. For collection, the cells were trypsinized, centrifuged and flash frozen. 50 million cells were collected for the final time point. The pooled competitive growth experiments were done in three (CALU1, NALM6, HAP1, K562, A172) or six (NCIH358, NCIH460, HCC827, HCT116) biological replicates.

gDNA from the cell samples was extracted with Qiagen Blood MIDI kit. The peptide expression cassette was PCR-amplified using Herculase II Fusion DNA Polymerase (600679, Agilent) from the gDNA pool in two steps. For PCR-1, twelve 100 µl reactions containing 10 µg gDNA each were performed to amplify the peptide cassettes together with adaptors for Illumina sequencing primers. The PCR-1 reactions from one sample were pooled and used as a template for PCR-2 to introduce the UDIs and Illumina adapters to the library. The PCR-2 was resolved on TAE agarose gels, extracted and purified with Monarch® DNA Gel Extraction Kit (New England Biolabs). The libraries were sequenced at Novogene on Novaseq X with paired-end 150 bp to a median depth of 475 reads per variant.

The paired-end reads were merged with FLASH^89^ version 1.2.11. The reads were mapped to the library variants using 2fast2q^90^. Peptide variants with less than 20 reads at t0 were dropped from further analysis. Then, the log2 fold change of peptide frequency in the sample collected after 8 cell doublings to the peptide frequency in the input sample (2 days after transduction) was calculated using the DESeq2 pipeline^91^. The cutoff for anti-proliferative peptides was set at log2 fold change -1 with multiple testing adjusted p-value cutoff at 0.0001.

### Recombinant peptide-GFP protein purification

For recombinant protein purification, the peptide-GFP constructs were cloned to pET28a, resulting in C-terminal tagging with 6xHis tag. The protein expression in *E. coli* BL21(DE3) cells was induced at culture optical density OD600=0.6 by addition of 1 mM IPTG, followed by further incubation at 37 °C for 3 h, after which the cells were pelleted and frozen. The cell pellets were resuspended in a buffer containing 50 mM Tris-HCl, pH 7.4, 300 mM NaCl, 1% Triton X-100, 5% glycerol, 10 mM imidazole and cOmplete, EDTA-free Protease Inhibitor Cocktail (Roche), and 1 mg/ml lysozyme (Sigma-Aldrich). The lysate was incubated on ice for 15 min, sonicated, and cleared by centrifugation at 4 °C 20 000 g for 15 min. The 6xHis-tagged proteins were bound to HisPur™ Ni-NTA Resin (Thermo Fisher Scientific) in gravity flow columns and washed 4 times with the lysis buffer supplemented with 25 mM imidazole. The proteins were eluted with 50 mM Tris-HCl, pH 7.4, 500 mM NaCl, 0.5% Triton X-100, 5% glycerol, and 200 mM imidazole. The eluates were analyzed by SDS-PAGE to verify the purity and to evaluate protein concentration using bovine serum albumin standards (Thermo Fisher Scientific).

### Peptide pulldown and HCF1 coIP

To identify the interactors of motif peptides, ChromoTek GFP-Trap® Magnetic Agarose (Proteintech) was used to pull down either peptide-GFP proteins expressed in cancer cell lines or to bind purified recombinant peptide-GFP proteins, followed by mixing with human cell lysate. For genetically expressed peptides, the peptide expression was induced with 1 µg/ml doxycycline at 40% confluency, the cells were harvested by scraping at 24 h and frozen. For pulldown experiments followed by immunoblotting, cells from one 10 cm dish were used, while cells from a 15 cm dish were used for mass-spectrometry experiments.

The cells were lysed in buffer containing 10 mM Tris-HCl, pH 7.4, 140 mM NaCl, 10% glycerol, 0.5% NP-40, 0.25% Triton X-100, 1 mM EDTA, 1 mM DTT and cOmplete^TM^ protein inhibitors cocktail (Merck). The lysates were pulled through a 26G needle and cleared by centrifugation at 20 000 g 4 °C 10 min. The total protein concentration in the lysate was measured using Pierce™ Bradford Plus Protein Assay Reagent (Thermo Fisher Scientific). The lysates were diluted to 2 mg/ml total protein. For co-immunoprecipitation, 1-2 mg of cell lysate was mixed with 10 µl (experiments analyzed by immunoblotting) or 20 µl (experiments analyzed by mass-spectrometry) ChromoTek GFP-Trap® Magnetic Agarose beads (Proteintech) that had been equilibrated with the lysis buffer. In experiments analyzed by mass-spectrometry, before mixing the lysate with GFP-trap beads, the lysate was precleared from non-specific binders using ChromoTek Binding Control Magnetic Agarose Beads (Proteintech). The mixture was incubated on a rotator at 4 °C for 1 hour. Then, the GFP-trap magnetic beads were collected using a magnet, the lysate was removed and the beads were washed three times with 1 ml lysis buffer. In case of pulldown experiments followed by mass-spectrometry, the beads were washed two more times with 100 mM ammonium bicarbonate, moving the beads to a new tube with each wash. For immunoblotting, the immunoprecipitated proteins were eluted using Laemmli SDS-PAGE sample buffer and were subjected to SDS-PAGE as above. Sample processing, including trypsinisation to peptides and peptide purification, and mass-spectrometry with Orbitrap Fusion Lumos Tribrid Mass Spectrometer (Thermo Fisher Scientific) was performed at the Cancer Research UK Cambridge Institute Proteomics core facility.

To characterize HCF1 interactome and its perturbations by the competitor peptides, HCT116 cells from 80% confluent 15 cm dish were harvested by scraping 24 h after inducing peptide-GFP expression with 1 µg/ml doxycycline. The cells were lysed as described above, followed by coimmunoprecipitation of HCF1 from HCT116 cell lysate with 1 µg anti-HCF1 (A301-400A, Bethyl Laboratories) bound to 25 µl protein A Dynabeads (10002D, Invitrogen). For negative control, rabbit IgG (#2729, Cell Signaling Technology) was used. The beads were washed, processed and analyzed for mass-spectrometry as above. Functional enrichment analysis of outcompeted proteins was performed with STRING^92^. For silver staining, the proteins were eluted using Laemmli sample buffer, resolved by SDS-PAGE and stained using SilverQuest™ Silver Staining Kit (Invitrogen).

### RNAseq

4×10^5^ NCIH358 cells transduced with peptide-GFP expression constructs were seeded to 6-well plate, cultured for 24 h, and induced for peptide expression with 1 µg/ml doxycycline. 24 h after induction, the cells were harvested by scraping and washed with PBS. RNA was extracted using High Pure RNA Isolation Kit (11828665001, Roche). All samples from one experiment were processed in parallel. Library preparation was performed at the Cancer Research UK Cambridge Institute Genomics core facility using Illumina Stranded mRNA Prep, followed by 2 x 50 paired-end sequencing on NovaSeq X (Illumina) to obtain at least 20 million reads per sample.

RNAseq data was processed with nf-core/rnaseq pipeline (version 3.17.0), using nf-core/differentialabundance pipeline^93^ to obtain genes that are differentially expressed with the peptide treatment. Functional enrichment analysis was performed with gProfiler^94^.

### qPCR

To investigate the effect of WRPW peptides and TLE1 overexpression on downstream gene expression, RNA was isolated with High Pure RNA Isolation Kit (11828665001, Roche) from 1 million NCIH358 cells harvested 24 h after induction of WRPW peptide expression with 1 µg/ml doxycycline. cDNA was synthesized with SuperScript™ IV Reverse Transcriptase (Invitrogen) using random hexamers as primers (N8080127, Invitrogen), and used as a template for qPCR with Luna® Universal qPCR Master Mix (New England Biolabs) in QuantStudio qPCR machine (Thermo Fisher Scientific). HPRT1 was used as a housekeeping gene. The qPCR experiments were done in biological duplicates with technical triplicates.

### Labelling of myristoylation with azido myristic acid

Azido myristic acid (HY-151855, Cambridge Bioscience) was dissolved at 50 mM in DMSO. HCT116 cells were seeded at 40% confluency in a 6-well plate, followed by peptide-GFP induction with addition of 1 µg/ml doxycycline. The culture medium was supplemented with 50 µM azido myristic acid. The cells were harvested 24h after treatment start. The cells were lysed and proteins extracted as described above (See Immunoblotting). ChromoTek GFP-Trap® Magnetic Agarose (Proteintech) beads were prepared by washing twice with cell lysis buffer. The azido myristic acid was labelled with TAMRA using the Click-iT Protein Reaction Buffer Kit (10005303, Thermo Fisher Scientific) as per manufacturer’s instructions using 15 µg total protein cell lysate and TAMRA-alkyne. Following the Click-IT reaction, the GFP-Trap beads were collected with a magnet, and the supernatant and bead-bound fraction were analyzed on SDS-PAGE. TAMRA fluorescence was detected using Li-Cor Odyssey M.

### Propidium iodide staining cell cycle analysis

HCT116 and NALM6 cells expressing cyclin D helical pocket competitor peptides were collected by trypsinization 16h after the start of peptide expression or treatment with palbociclib. The harvested cells were washed with PBS, pelleted, and fixed in -20 °C 70% ethanol at 4 °C for at least 30 min. The ethanol-fixed cells were washed twice with PBS and resuspended in 300 µl solution containing 5 µg/ml propidium iodide and 25 µg/ml RNase A in PBS. The mixture was incubated at 37 °C in the dark for 30 minutes and analyzed by flow cytometry using BD LSR Fortessa (BD Biosciences). Data from at least 10 000 cells was collected and analyzed using FlowJo.

### Microscopy

HCT116 cells transduced with HES1 WRPW-GFP expression constructs were grown to 30% confluency in a Cellvis P24-1.5P 24-well plate with polymer coverslip base (CellVis), followed by induction of peptide expression with 1 µg/ml doxycycline for 24h. The cells were stained for 10 min with 8 µM Hoescht-33342 (TargetMol), followed by washing twice with medium and imaging with Operetta CLS (Revvity) microscope using 40x objective.

### Cell cycle arrest Incucyte experiments

The FUCCI G1-phase reporter mCherry-CDT1^95^ was expressed as a T2A fusion to peptide-GFP to monitor the cell cycle phase in GFP-expressing cells. Cells were cultured to be at 50% confluency before setting up for time-lapse fluorescence imaging with Incucyte S3 (Sartorius). 5,000 or 15,000 cells were seeded in a 96-well plate for adherent or suspension cell lines, respectively. The mCherry-CDT1 and peptide-GFP expression was induced with 1 µg/ml doxycycline. Palbociclib concentrations ranged from 0.3 to 10,000 nM. The cells were imaged with phase-contrast, GFP (300 ms), and mCherry (400 ms) every 3 hours using a 10x objective with S3/SX1 G/R Optical Module. The cells were segmented using Adherent Cell-by-Cell segmentation and the GFP-positive cells were divided based on high or low mCherry signal.

### Cyclin-CDK and RbC protein expression and purification

Human cyclin–Cdk fusion complexes were expressed in *Saccharomyces cerevisiae* and purified as previously described^96,97^. Briefly, pRS425-pGAL1-3×FLAG–cyclin D–L–Cdk6 plasmids (where L denotes a 3×GGGGS linker) were transformed into yeast, and protein expression was induced at an OD_₆₀₀_ of 0.6–0.8 for 3 h at 30 °C. Following harvesting and lysis, cyclin-Cdk complexes were isolated using anti-FLAG M2 affinity agarose beads (Sigma-Aldrich, #A2220) before being eluted in buffer containing 50 mM HEPES-KOH (pH 7.6), 250 mM KCl, 1 mM MgCl_₂_, 1 mM EGTA, 5% glycerol, and 0.2 mg/ml 3×FLAG peptide (Sigma-Aldrich, #F4799). Purified complexes were verified via western blot using mouse anti-FLAG antibody (1:2,000; Sigma-Aldrich, #F1804) coupled with an anti-mouse IRDye 800CW secondary antibody (1:10,000; LI-COR, #926-32212).

An N-terminally His-tagged C-terminal fragment of RB1 (RbC; RB1 amino acids 772–928) was expressed in *Escherichia coli* and purified by cobalt affinity chromatography. Specifically, the pET28a-6×His-RbC plasmid was transformed into *E. coli* BL21(DE3)RIL cells (Agilent, #230245). Protein expression was induced with 1 mM IPTG, and cultures were incubated at 30 °C for 3 h. Cells were harvested by centrifugation and lysed in a B-PER™ Complete Bacterial Protein Extraction Reagent (ThermoFisher # 89821) supplemented with protease inhibitors for 15 minutes at room temperature. Lysates were diluted with a wash buffer (25 mM HEPES, pH 7.4; 300 mM NaCl; 10% glycerol) and clarified by centrifugation. The His-tagged protein was purified by gravity-flow chromatography using cobalt affinity resin (HisPur™ Cobalt Resin, Thermo Scientific, #89965). The resin was washed sequentially with 20 resin volumes, followed by 10 resin volumes of wash buffer containing 10 mM imidazole then 10 column volumes of wash buffer containing 20 mM imidazole. Bound protein was eluted in a wash buffer supplemented with 200 mM imidazole.

### Synthetic peptides

Peptides were synthesized by Peptide 2.0 Inc. Lyophilized peptides were resuspended in 50 mM HEPES pH 7.4. The RBL2 helix peptide sequence is SALLRRLQDVANDRGS.

### In vitro kinase assays with peptide competition

In vitro kinase assays using ATP-γ-S were performed as previously described97, with minor modifications. Reactions were carried out in 50 mM HEPES (pH 7.4), 150 mM NaCl, 5 mM MgCl_₂_, 0.5 mM DTT, 0.08 mg/ml BSA, 2% DMSO, and 0.5 mM ATP-γ-S. FLAG elution buffer was added as required to normalize reaction volumes across different cyclin–CDK complexes. Enzyme concentrations were kept in the low nanomolar range and adjusted such that less than 10% of substrate was converted over the course of the reaction, unless stated otherwise. Competitor peptide concentration series were designed to span the full IC_₅₀_ range. Reaction mixtures containing 1 μM RbC, cyclin-CDK complexes, and helix peptides at the indicated concentrations were allowed to equilibrate for ≥ 8 minutes before reactions were initiated by the addition of ATP-γ-S. Reactions were incubated at room temperature for 12 min and quenched by addition of EDTA to a final concentration of 40 mM. Thiophosphorylated substrates were alkylated with 2.5 mM p-nitrobenzyl mesylate (pNBM; Selleckchem, #E1248) for 1 h at room temperature. Alkylation was terminated by addition of 20 mM DTT and Laemmli sample buffer (Bio-Rad, #1610747), followed by heating at 70 °C for 10 min and centrifugation at 16,000 g for 1 min. Reactions were resolved via SDS-PAGE using 12% acrylamide gels and phosphorylated RbC was detected via western blot using a rabbit anti-thiophosphate ester antibody (1:2000, Abcam, ab133473) coupled with coupled with an anti-rabbit IRDye 800CW secondary antibody (1:12,000; LI-COR, #926-32213). Kinase activity was quantified using Image Studio v6.0 (LI-COR) and plotted using Prism v10.0 (Graphpad).

