## Supplementary Figure 1 for "Uncovering cancer dependencies in peptide-interacting protein pockets"

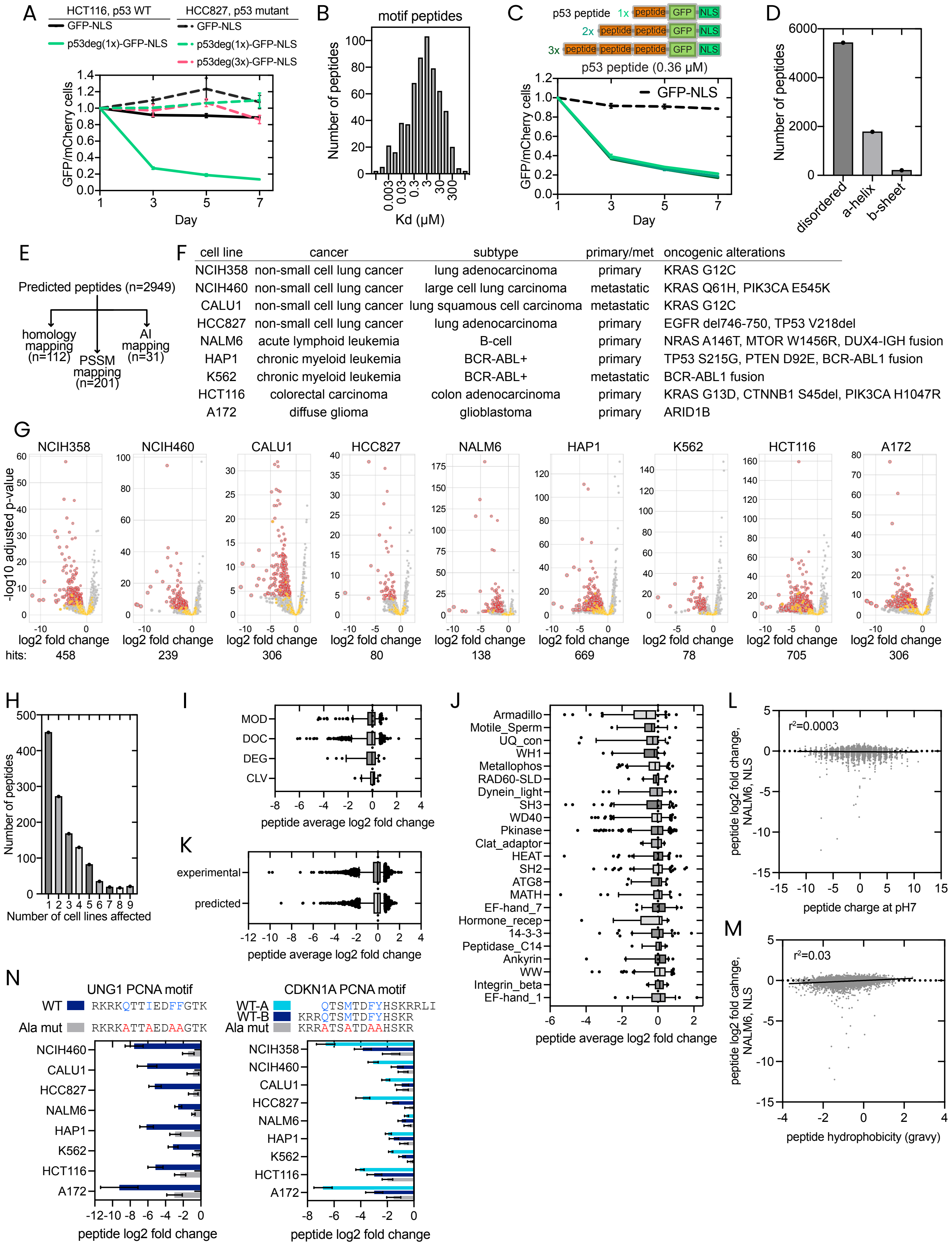


| cell line | cancer | subtype | primary/met | oncogenic alterations |
|---|---|---|---|---|
| NCIH358 | non-small cell lung cancer | lung adenocarcinoma | primary | KRAS G12C |
| NCIH460 | non-small cell lung cancer | large cell lung carcinoma | metastatic | KRAS Q61H, PIK3CA E545K |
| CALU1 | non-small cell lung cancer | lung squamous cell carcinoma | metastatic | KRAS G12C |
| HCC827 | non-small cell lung cancer | lung adenocarcinoma | primary | EGFR del746-750, TP53 V218del |
| NALM6 | acute lymphoid leukemia | B-cell | primary | NRAS A146T, MTOR W1456R, DUX4-IGH fusion |
| HAP1 | chronic myeloid leukemia | BCR-ABL+ | primary | TP53 S215G, PTEN D92E, BCR-ABL1 fusion |
| K562 | chronic myeloid leukemia | BCR-ABL+ | metastatic | BCR-ABL1 fusion |
| HCT116 | colorectal carcinoma | colon adenocarcinoma | primary | KRAS G13D, CTNNB1 S45del, PIK3CA H1047R |
| A172 | diffuse glioma | glioblastoma | primary | ARID1B |

**Figure S1. Peptide expression optimisation, proteome-wide motif competitor library design and screening.** (**A**) Pairwise competitive growth experiment results showing that the competitor peptides containing the p53 degron are anti-proliferative in p53-WT HCT116 cells, but not in p53 mutant HCC827 cells. (**B**) Distribution of measured dissociation constants (Kd) of motif-mediated interactions from the ELM database16. (**C**) Pairwise competitive growth experiments showing the impact of motif repeats on the potency of competitors based on p53 peptide. (**D**) The AlphaFold2-predicted[85] secondary structure of the 15mer peptides in the proteome-wide motif peptide library. (**E**) The predicted motif peptides were subjected to motif mapping based on homology, PSSM scoring and AI to predict motif classes for 344 peptides. (**F**) The characteristics of cell lines used in the proteome-wide competitor peptide dropout screen. (**G**) Vulcano plots showing the effect of peptides on cell proliferation in the proteome-wide peptide dropout screen. (G) The number of cell lines affected by the detected anti-proliferative peptides. (**H**) Distribution of peptide dropout scores based on their ELM class (MOD - modification; DOC - docking; DEG - degron; CLV - cleavage). The plot contains averaged data from all cell lines. (**I**) Distribution of dropout scores of peptides binding to proteins of different domain families. The plot contains averaged data from all cell lines. (**J**) Experimentally characterised and predicted motif peptide dropout scores. The plot contains averaged data from all cell lines. (**L**,**M**) Plots showing the peptide dropout scores together with charge (L) or hydrophobicity (M). (**N**) Peptide dropout scores from the proteome-wide screen for wild-type and alanine mutant PCNA-binding peptides. Error bars show 95% confidence intervals of the median.
