## Supplementary Figure 2 for "Uncovering cancer dependencies in peptide-interacting protein pockets"

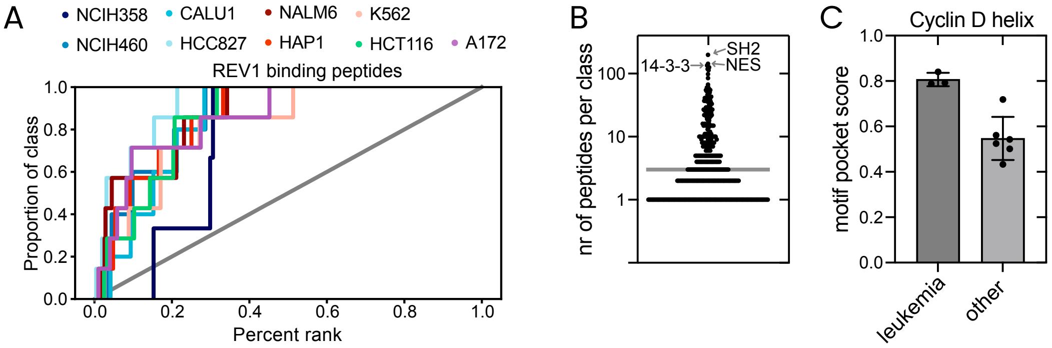


D

| essentiality | | | | | | |
|---|---|---|---|---|---|---|
| pocket | gene | proteins | paralogs | % | syn. let. | % |
| + | - | 76 | 29 | 38% | 6 | 21% |
| + | + | 37 | 11 | 30% | 3 | 27% |
| - | - | 149 | 79 | 53% | 5 | 6% |
| - | + | 18 | 1 | 6% | 0 | 0% |

paralog pairs

- BRD2-BRD3
- CCND1-CCND3
- CTBP1-CTBP2
- KAT2A-KAT2B
- KEAP1-KLHL2
- STAG1-STAG2

**Figure S2. Proteome-wide characterisation of competitor peptide sensitivities.** (**A**) Cumulative dropout rank plots of REV1_RIR peptides in 9 cancer cell lines in the proteome-wide peptide competitive growth experiment. (**B**) Distribution of the number of peptides in the library belonging to different motif families. The line shows the median. (**C**) Compared to adherent cell lines (NCIH358, NCIH460, CALU1, HCC827, HCT116, A172), the three studied leukemia cell lines (NALM6, HAP1, K562) have higher sensitivity to peptides that bind to the cyclin D pocket that recruits RB1 and RBL1/2. (**D**) Table showing the classification of the target pocket dependencies in Fig. 2D, and the number and percentage of paralogs and synthetic lethal pairs in different categories.
