## Supplementary Figure 3 for "Uncovering cancer dependencies in peptide-interacting protein pockets"

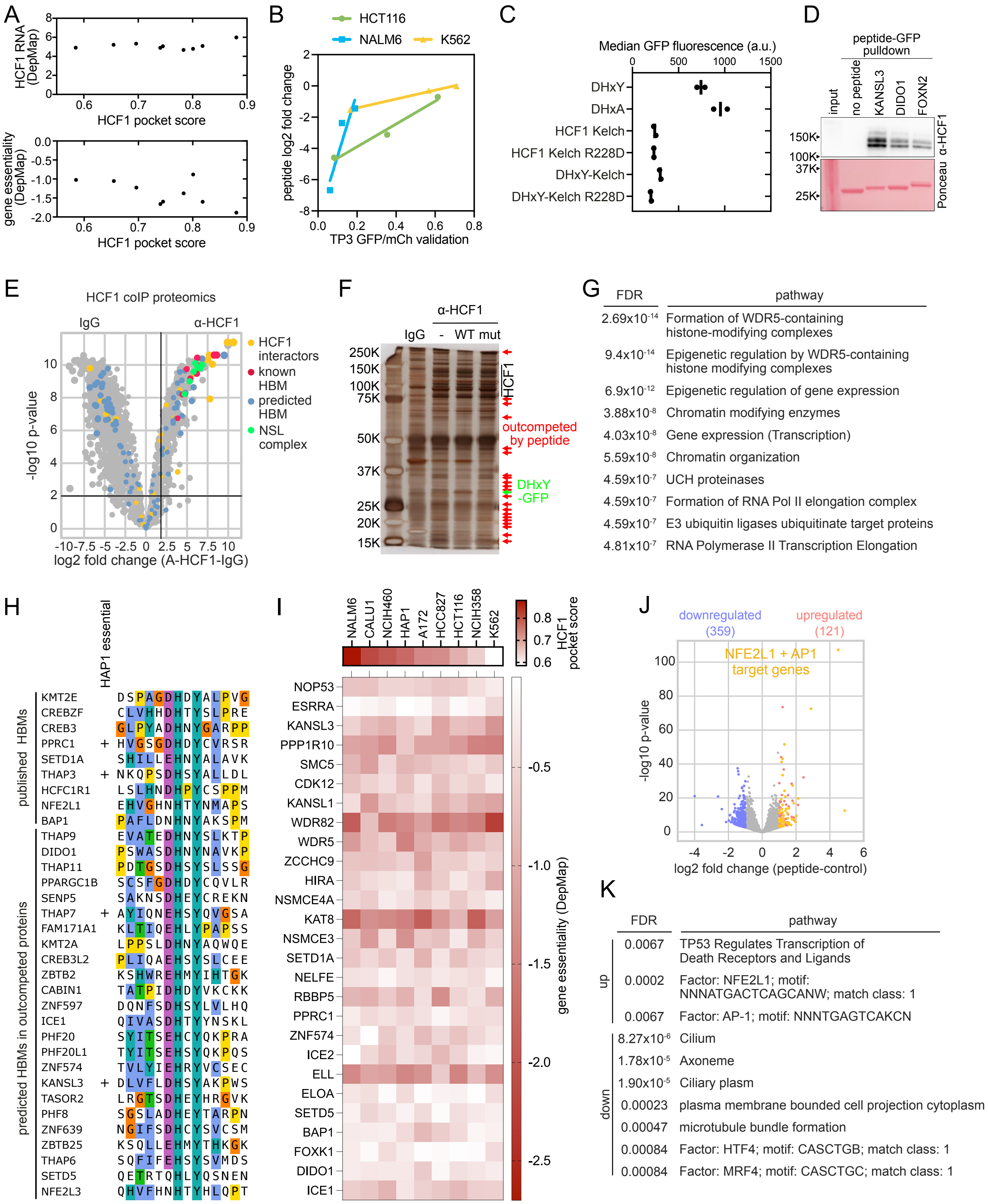

**Figure S3. Characterisation of HCF1-targeting competitor peptide phenotype.** (**A**) Correlation between the HCF1 motif pocket score obtained from the proteome-wide screen and HCF1 expression (RNAseq data from DepMap, top) or HCF1 gene essentiality (DepMap CRISPR knockout score, bottom). (**B**) Correlation between the proteome-wide screen dropout scores and the anti-proliferative effect in validation GFP/mCherry pairwise competitive growth experiments for HCF1-targeting peptides from KANSL3, DIDO1 and FOXN2. (**C**) GFP fluorescence intensities of cells expressing the constructs in Fig. 3J shows the lower expression of the Kelch domain compared to the peptides. (**D**) Pulldown of HCF1 from HCT116 lysate using recombinant peptide-GFP proteins. A representative example from three biological replicates is shown. (**E**) HCF1 coIP from HCT116 cell lysate followed by mass-spectrometry to detect HCF1 interactors. (**F**) Silver-stained SDS-PAGE of HCF1 coIP from HCT116 cell lysate in the presence or absence of the KANSL3 HCF1 competitor peptide reveals that the peptide decreases the abundance of many HCF1 interactors. (**G**) GO term enrichment analysis of the HCF1 interactors that are outcompeted by the KANSL3 peptide. (**H**) Alignment of the validated and potential HCF1 binding motifs in the HCF1 interactors that are outcompeted by the competitor peptide in Fig. 3K. Four of these motifs were identified as essential motifs in a base editing screen in HAP18. (**I**) Heatmap showing the DepMap gene dependency scores of the proteins outcompeted from HCF1 binding by the KANSL3 peptide. Only essential genes (minimal score <-0.5) are shown. (**J**) Up- and down-regulated genes detected by RNAseq in HCT116 cells treated with KANSL3 peptide for 24h. (**K**) The top enriched GO terms from the up- and downregulated genes in RNAseq data of HCT116 cells overexpressing the KANSL3 peptide.
