## Supplementary Figure 4 for "Uncovering cancer dependencies in peptide-interacting protein pockets"

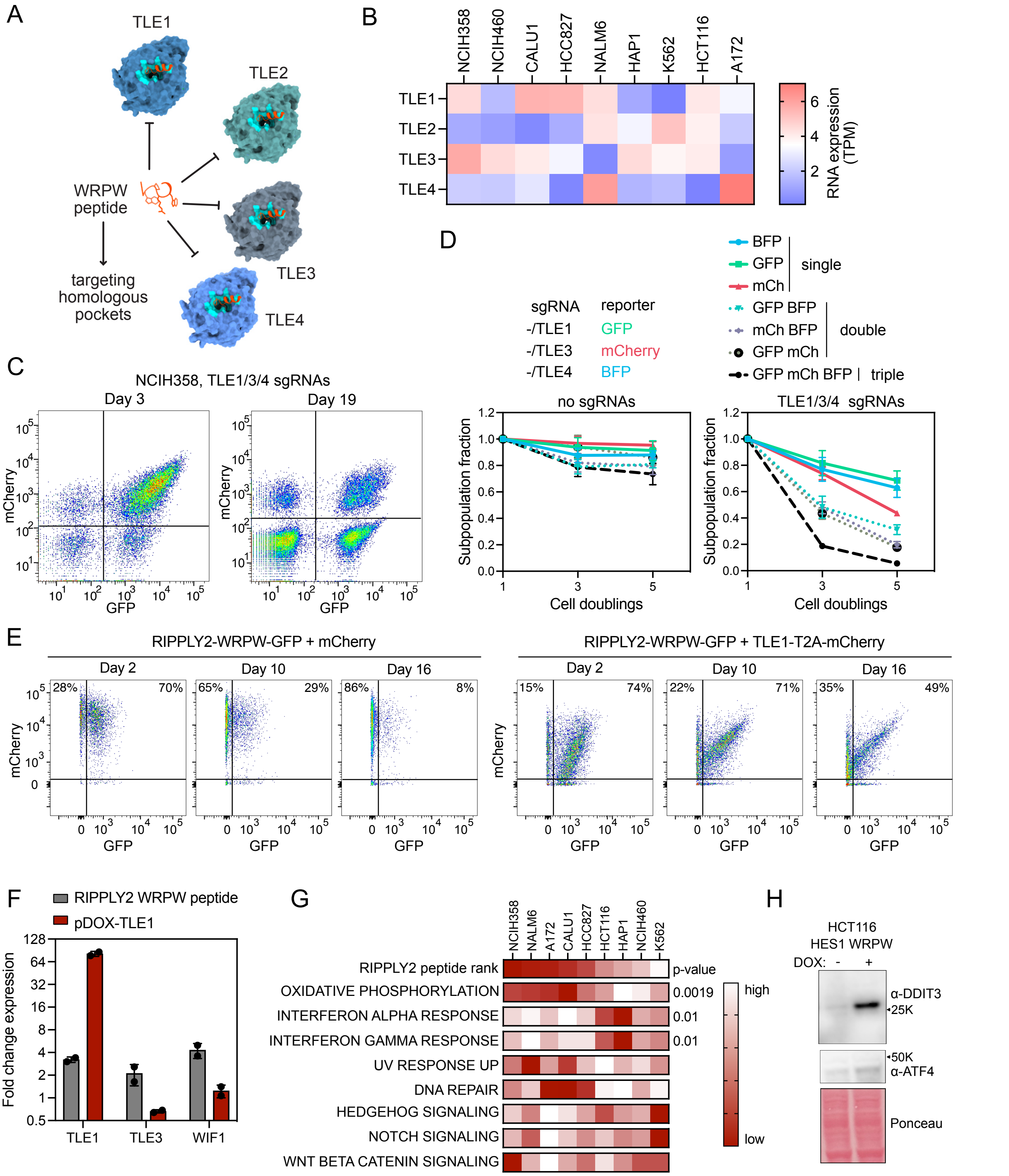


**Figure S4. Characterisation of TLE knockout and overexpression phenotypes.** (**A**) Schematic model of a WRPW peptide targeting homologous pockets in TLE1-4 WD40 domains. TLE1-4 in complex with WRPW peptide was modelled with AlphaFold3. (**B**) Gene expression of TLE1/2/3/4 in the indicated cell lines. RNAseq data from DepMap. (**C**) Example of monitoring subpopulations of NCIH358 cells expressing different gRNAs and fluorescent reporters in a pooled multiplex TLE1/3/4 CRISPR knockout competitive growth experiment. (**D**) Time-course of relative fractions of different NCIH358 cell subpopulations expressing the indicated gRNAs and fluorescent reporters measured by flow cytometry. At each time point, the culture was split ¼ and cultured to 80% confluency. (**E**) Example of gating for flow-cytometry-based competitive growth experiment presented in Fig. 4F. (**F**) The effect of TLE1 or RIPPLY2 WRPW peptide overexpression on TLE1, TLE3 and WIF1 gene expression measured by qPCR. (**G**) Hallmark 50 gene set activities were estimated from RNAseq data using ssgsea. T-test was used to compare pathway activities in RIPPLY2-sensitive (NCIH358, NALM6, CALU1, A172) cell lines and -insensitive lines (HCT116, NCIH460, K562, HAP1). Heatmaps showing the RIPPLY2 peptide rank and scores for selected gene sets in the 9 cancer cell lines. (**H**) Western blot showing ATF4 response upon HES1 WRPW peptide expression in HCT116 cells. A representative example from two biological replicates is shown.
