## Supplementary Figure 5 for "Uncovering cancer dependencies in peptide-interacting protein pockets"

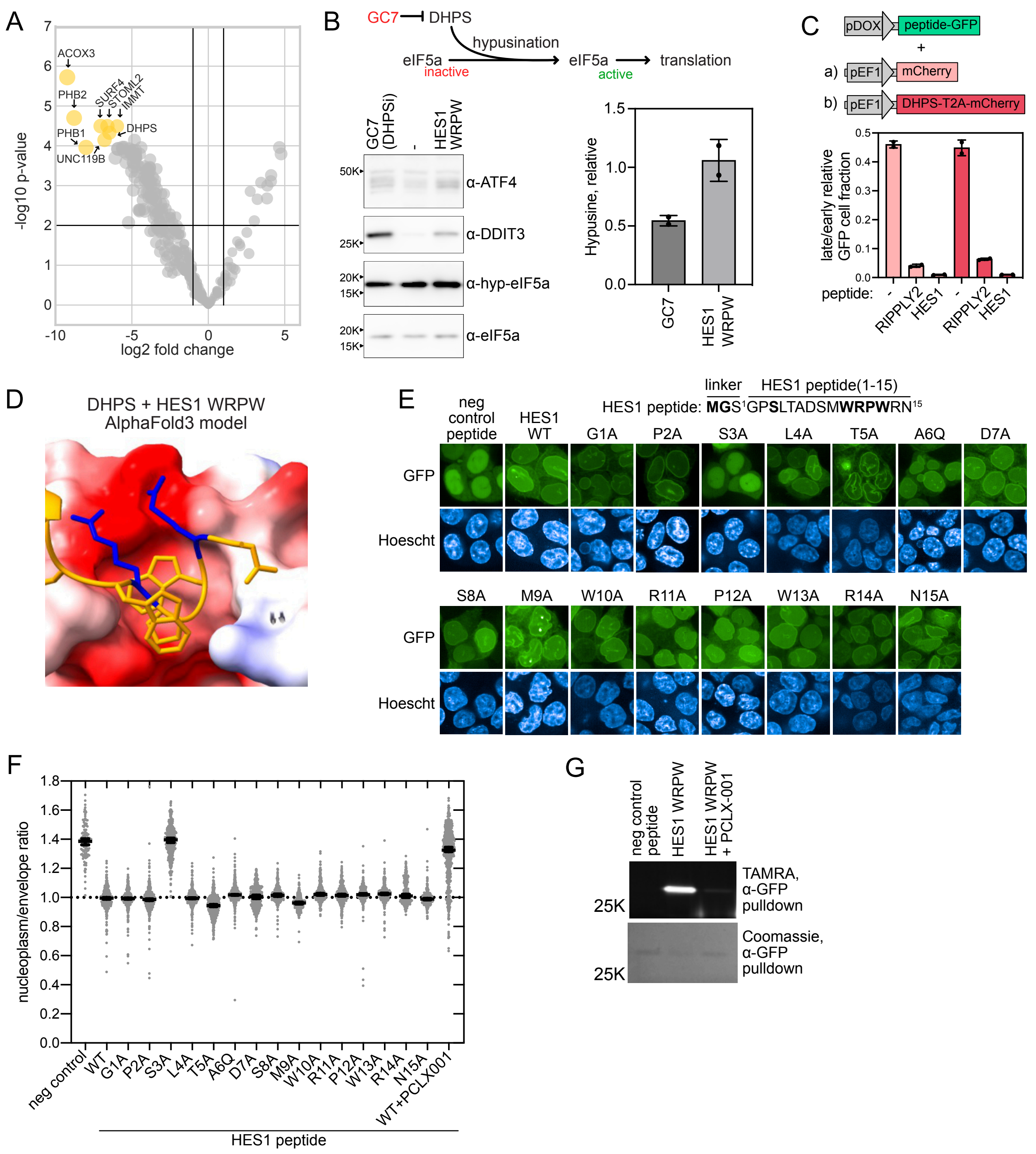


**Figure S5. Characterisation of HES1 WRPW peptide phenotype.** (**A**) Mass-spectrometry data comparing proteins pulled down by RIPPLY2 and HES1 WRPW peptide-GFP fusions from NCIH358 lysates. (**B**) Western blots showing the effect of GC7, a small molecule inhibitor of DHPS and HES1 WRPW peptide on ATF4 response and hypusination of eIF5a in NCIH358 cells treated for 48h. (**C**) The impact of DHPS-T2A-mCherry overexpression on the proliferation rate of cells expressing RIPPLY2 and HES1 WRPW GFP fusions in a flow-cytometry-based competitive growth experiment. Mean±standard deviation from two biological replicates is shown. (**D**) AlphaFold3 structural model of HES1 WRPW bound to DHPS, shown in electrostatic coloring, with two Arg residues unique to HES1 among the WRPW peptides binding to an acidic patch in DHPS. (**E**) Microscopy images showing subcellular localisation of HES1 WRPW peptide single mutants in HCT116 cells. (**F**) Ratio of peptide-GFP fluorescence signal in the nucleoplasm to the nuclear envelope in different HES1 WRPW peptide variants. Each data point is data from a single cell. Bars: median±95% confidence intervals. (**G**) Myristoylation of HES1 peptide in untreated and PCLX-001-treated HCT116 cells using TAMRA-alkyne to detect azido-myristate.
