## Supplementary Figure 6 for "Uncovering cancer dependencies in peptide-interacting protein pockets"

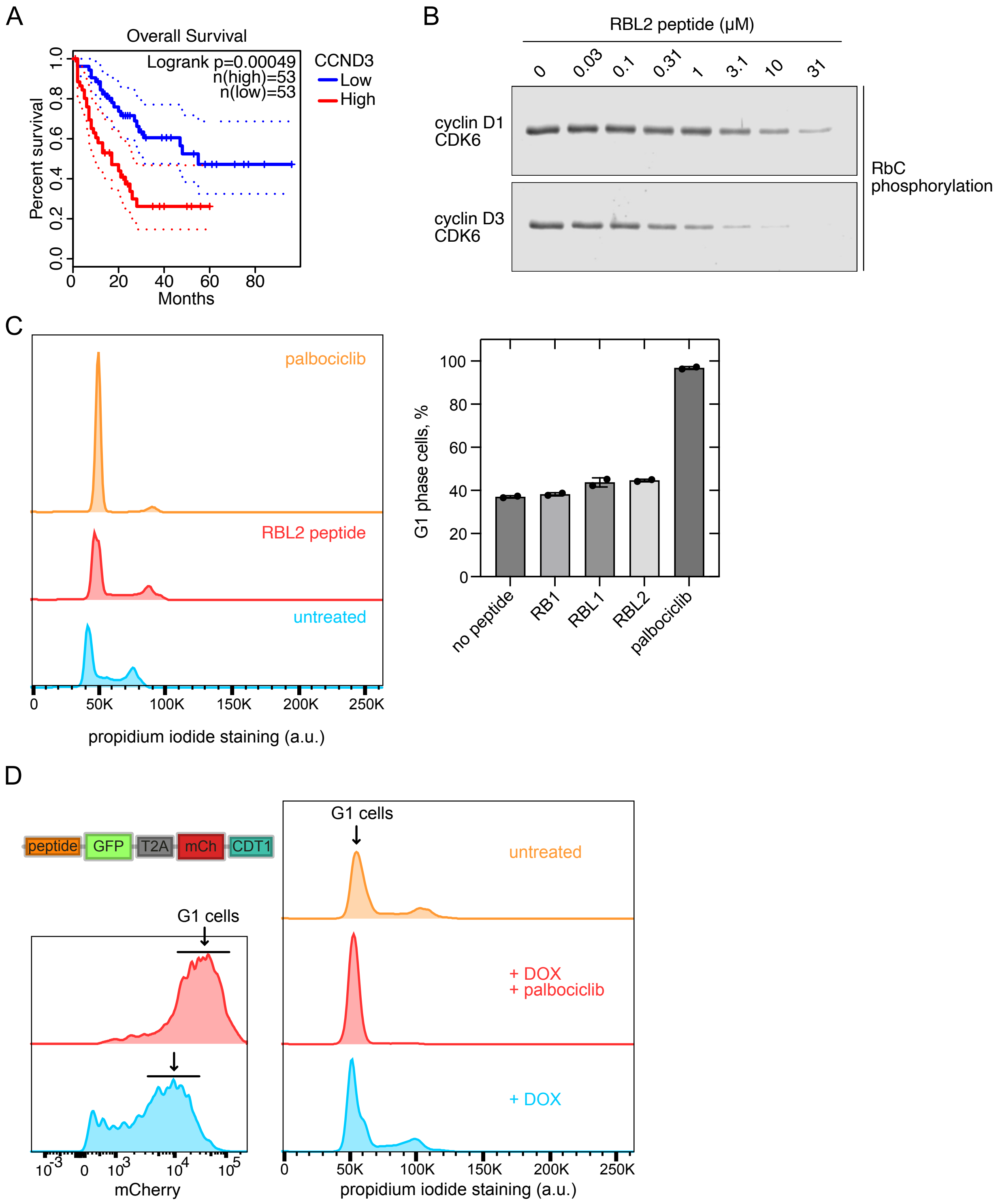


**Figure S6. The impact of cyclin D3 in AML patients and cell lines, and validation of mCherry-CDT1 G1-phase reporter.** (**A**) High cyclin D3 expression is associated with poor prognosis in acute myeloid leukaemia patients (TCGA data). (**B**) In vitro RbC phosphorylation assay measuring the dose-sensitivity of cyclin D1-CDK6 and cyclin D3-CDK6 to RBL2 peptide. A representative example of three experiments is shown. (**C**) The effect of cyclin D targeting peptide expression on increasing the G1-phase cell fraction in NALM6 cell populations expressing the indicated peptides or treated with 300 nM palbociclib. DNA was stained with propidium iodide and analyzed by flow cytometry. On the left, a representative experiment is shown. (**D**) mCherry-CDT1 sensor was validated for G1-phase quantification by comparison with cell cycle phase distribution measured by DNA staining with propidium iodide staining.
