## Supplementary Figure 7 for "Uncovering cancer dependencies in peptide-interacting protein pockets"

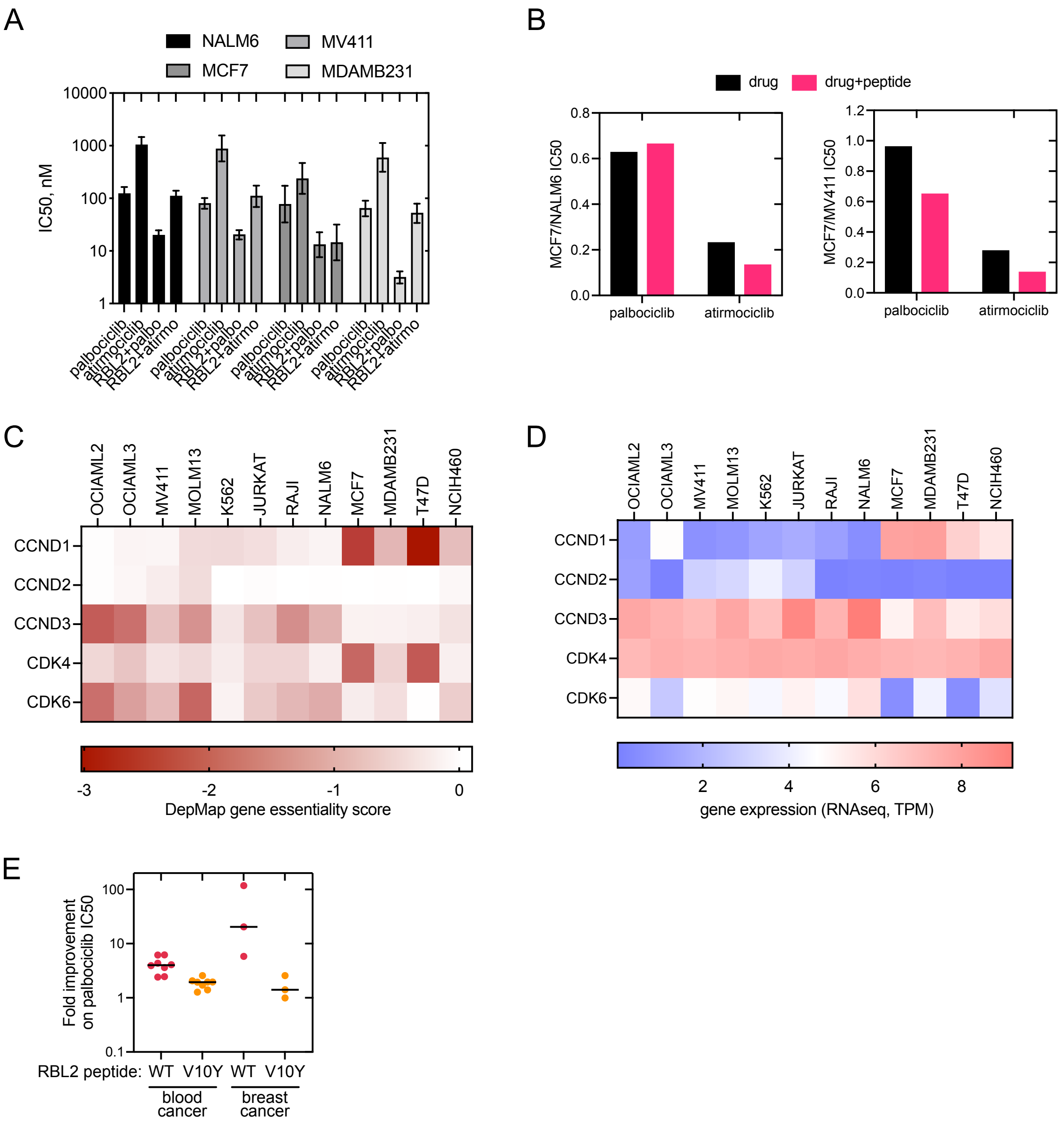


**Figure S7. The impact of cyclin specificity of cyclin D targeting peptides on G1-phase.** (**A**) The IC50 of palbociclib and atirmociclib in the presence or absence of the RBL2 competitor peptide measured by the G1 cell cycle phase fraction 24h after treatment start. (**B**) The relative IC50 values of ATP-competitive CDK4/6 inhibitors in the presence or absence of RBL2 competitor peptide in MCF7 cells compared to NALM6 (left) and MV411 (right) cells. (**C**) DepMap gene essentiality scores of cyclin D1-D3 and CDK4/6 in different cancer cell lines. (**D**) RNA expression scores (counts per million) of cyclin D1-D3 and CDK4/6 in different cancer cell lines. Data is from DepMap. (**E**) The fold improvement of palbociclib IC50 in the presence of wild-type or V10Y mutant RBL2 peptide in blood (OCIAML2, OCIAML3, MV411, MOLM13, K562, JURKAT, RAJI, NALM6) and breast cancer cell lines (MCF7, MDA-MB-231, T47D). Palbociclib IC50 was measured at 24h after treatment using CDT1-mCherry as a reporter for G1 phase cell cycle arrest.
